# NF-κB restrains nutrient-dependent transcription programs through chromatin modulation in *Drosophila*

**DOI:** 10.1101/2025.10.24.684223

**Authors:** Xiangshuo Kong, Conghui Li, Jason Karpac

**Affiliations:** Department of Biology, Texas A&M University, College Station, TX, 77843, USA; Department of Cell Biology and Genetics, College of Medicine, Texas A&M University, Bryan, TX, 77807, USA

## Abstract

The co-evolution of immune and metabolic systems has endowed immune signaling pathways with distinct control of cellular metabolism. Innate immune transcription factors, such as nuclear factor κB (NF-κB), have thus emerged as key regulators of adaptive metabolic responses to changes in diet and nutrition. Utilizing chromatin accessibility genomics, we found that Drosophila NF-κB (Relish) can restrain nutrient-dependent metabolic transcriptional programs that control cellular catabolism of energy substrates, divergent from the protein’s canonical role as a transcriptional activator. NF-κB/Relish restricts chromatin accessibility through modulating histone acetylation at metabolic target gene loci, which restrains metabolic gene transcription and blocks excessive activation of nutrient-dependent metabolic programs. Targeted genetic screening revealed that histone deacetylase 6 (HDAC6) interacts with NF-κB/Relish at NF-κB DNA regulatory motifs to limit chromatin accessibility and repress metabolic transcriptional programs. These results highlight that innate immune transcription factors can epigenetically restrain cellular catabolism to fine-tune nutrient-dependent metabolic adaptation.

---

**Introduction:** Mounting an immune response to a pathogenic challenge is energy-intensive and represents an inherent energetic trade-off with other physiological processes^1–6^. As a result, immune and metabolic systems are evolutionarily coordinated, reflecting an ancient co-adaptation to the energetic demands of host defense. Conversely, when homeostatic energy balance is disrupted by nutrient deprivation or nutrient surplus (often through changes in key energy substrates such as amino acids, sugars, and lipids), immune function is frequently compromised^1, 2, 7, 8^. The integration of immune and metabolic signaling architecture is thus essential to adjust pathogen-dependent and/or nutrient-dependent physiology and pathophysiology in animals across taxa, including humans.

Among immune regulators involved in immune-metabolic crosstalk, nuclear factor kappa-light-chain-enhancer of activated B cells (NF-κB) transcription factors serve as evolutionarily conserved central nodes in signaling integration related to metabolism, innate immunity, and adaptive immunity^9, 10^. Studies have also revealed an intimate relationship between NF-κB signaling axes and distinct aspects of metabolic regulation^11–14^. This includes, for example, the ability of NF-κB to dictate transcriptional programs and shape cellular energy stores by influencing lipid turnover through lipolysis^15, 16^. NF-κB activation in a variety of vertebrate cell types has been shown to promote triacylglycerol (stored lipid) catabolism through elevating CES1 lipase gene expression to mobilize lipid storage for energy provision during nutrient deprivation^15^. Also, in invertebrates, we previously reported that NF-κB transcription factors play a critical role in the metabolic adaptation to nutrient-deprivation by suppressing triacylglycerol lipolysis through transcriptional attenuation of lipase gene expression^16^. These findings and others demonstrate that the NF-κB transcriptional machinery is intricately (and evolutionarily) wired into a wide range of tissue- and context-specific metabolic programs. Thus, dissecting the mechanistic integration between NF-κB and distinct metabolic networks is essential for understanding its role in the regulation of metabolism in response to dietary, nutrient, or pathogenic challenges.

Although NF-κB transcription factors are classically recognized as activators of gene expression, accumulating evidence suggests that they can also exert context-dependent (and maybe signaling pathway distinct) transcriptional repression or attenuation. Several mechanisms have been proposed to explain this repressive function, including competition with other transcriptional activators for DNA binding^17, 18^, recruitment of corepressor complexes such as histone deacetylases (HDACs)^19–21^, induction of repressive complexes that interfere with coactivator function^22, 23^, post-transcriptional gene silencing^24^, and indirect suppression via feedback regulators^25, 26^. Just like immune transcriptional programs, the regulation of metabolic transcriptional networks must be fine-tuned to balance inducibility with transcriptional repression to maintain physiology long-term, such as sustaining energy substrate storage and usage. For example, lipids (especially triglycerides) represent an energetically concentrated way of storing energy across the evolutionary spectrum, and many eukaryotes are furnished with robust molecular machinery to convert other nutrients, such as amino acids and sugars, into stored lipids^27, 28^. Thus, lipid catabolic and anabolic transcriptional networks are balanced by activating and repressive mechanisms. Previous studies suggest that NF-κB transcription factors play a role in tuning these transcriptional networks, including through restraining gene expression, to direct nutrient-dependent metabolic programs^16^.

Adaptive metabolic mechanisms have evolved under intense selective pressure from both food scarcity and pathogen density. To this end, NF-κB-dependent control of metabolic programs presumably evolved, in part, from ancient responses to nutrient deprivation (or fasting), while some of these mechanisms are likely repurposed to control adaptive energy substrate storage in response to nutrient imbalances, such as overnutrition in humans. Fasting induces rapid and coordinated transcriptional activation of genes involved in nutrient mobilization, energy production, and stress adaptation^29–31^. Increasingly, these transcriptional responses have been shown to depend on chromatin remodeling, including dynamic changes in histone modifications, nucleosome positioning, and chromatin accessibility^32–34^. Such epigenomic regulation allows tight temporal and spatial control of metabolic gene expression, balancing transcriptional activation and repression. Notably, emerging studies indicate that NF-κB may directly or indirectly influence chromatin organization^35–37^. Understanding how this central immune-metabolic regulator coordinates nutrient signaling and epigenomic outputs may provide critical insights into the transcriptional and chromatin-level coordination of metabolic adaptation.

Invertebrate models hold distinct advantages to dissecting the integration of immune-metabolic signaling architecture and ancient immune transcription factor functions. These include the convergence of immune and metabolic functions within isolated tissues, unlike the spatial and functional separation of immune and metabolic organs in vertebrates^38–42^. Here, we exploited the fruit fly *Drosophila melanogaster* and uncovered that a Drosophila NF-κB transcription factor (Relish) plays a critical role in safeguarding metabolic homeostasis during nutrient deprivation, restraining fasting-induced chromatin changes and limiting transcriptional activation of catabolic gene networks. By integrating chromatin accessibility genomics, genetic screening, and molecular biochemistry, we identified HDAC6 as a key co-regulator that interacts with NF-κB/Relish to limit metabolic gene transcriptional induction during fasting. These findings uncover a previously unrecognized immune-metabolic integrative mechanism by which NF-κB cooperates with HDAC6 to fine-tune chromatin dynamics and transcriptional outputs, thereby preventing metabolic hyperactivation and preserving fitness during metabolic adaptation to nutrient challenges.

## Results

### NF-κB/Relish regulates nutrient-dependent chromatin remodeling

To comprehensively explore the genome-wide role of NF-κB in shaping transcriptional responses to metabolic adaptation, we first employed ATAC-seq to profile chromatin accessibility changes during acute fasting (before mortality) in Drosophila lacking functional NF-κB/Relish (utilizing the *rel^E^*^20^ allele). Relish is similar to mammalian p100/p105 NF-κB proteins and contains a Rel-homology domain, as well as ankyrin repeats (found in mammalian inhibitory IκBs)^43–45^. Principal component analysis (PCA) revealed distinct chromatin accessibility landscapes between wild-type control OreR and *rel^E^*^20^ mutant adult female flies (Figure S1A), independent of nutritional conditions, indicating an intrinsic role of NF-κB/Relish in modulating chromatin accessibility. Notably, the *Relish* gene locus itself showed significantly reduced chromatin accessibility in *rel^E^*^20^ mutants compared to controls (Figure S1B), corresponding to the known deletion of this region in the mutant allele. To further characterize genomic functional elements, we used HOMER to annotate the genomic features of accessible regions, and we found that nutrient deprivation did not alter the composition of functional elements within accessible regions in controls (Figure 1A). In contrast, NF-κB/Relish mutant flies exhibited a marked redistribution of chromatin accessibility upon fasting, with a substantial proportion of accessible regions enriched at promoter-TSS elements, but also within intronic and exonic genomic loci (Figure 1A). This genome-wide alteration of functional regions indicated that NF-κB/Relish may regulate the expression of a broad set of genes in response to fasting. Additionally, over 5,000 ATAC-seq peaks gained accessibility in *rel^E^*^20^ mutants after starvation and only 132 peaks gained accessibility in OreR flies (Figure 1B). Heatmap analysis confirmed that these differentially accessible peaks were highly consistent within each group (Figure 1C). These findings suggest that fasting induces widespread chromatin remodeling in the absence of NF-κB, relevant for the regulation of gene expression during metabolic adaptation.

**Figure 1.**
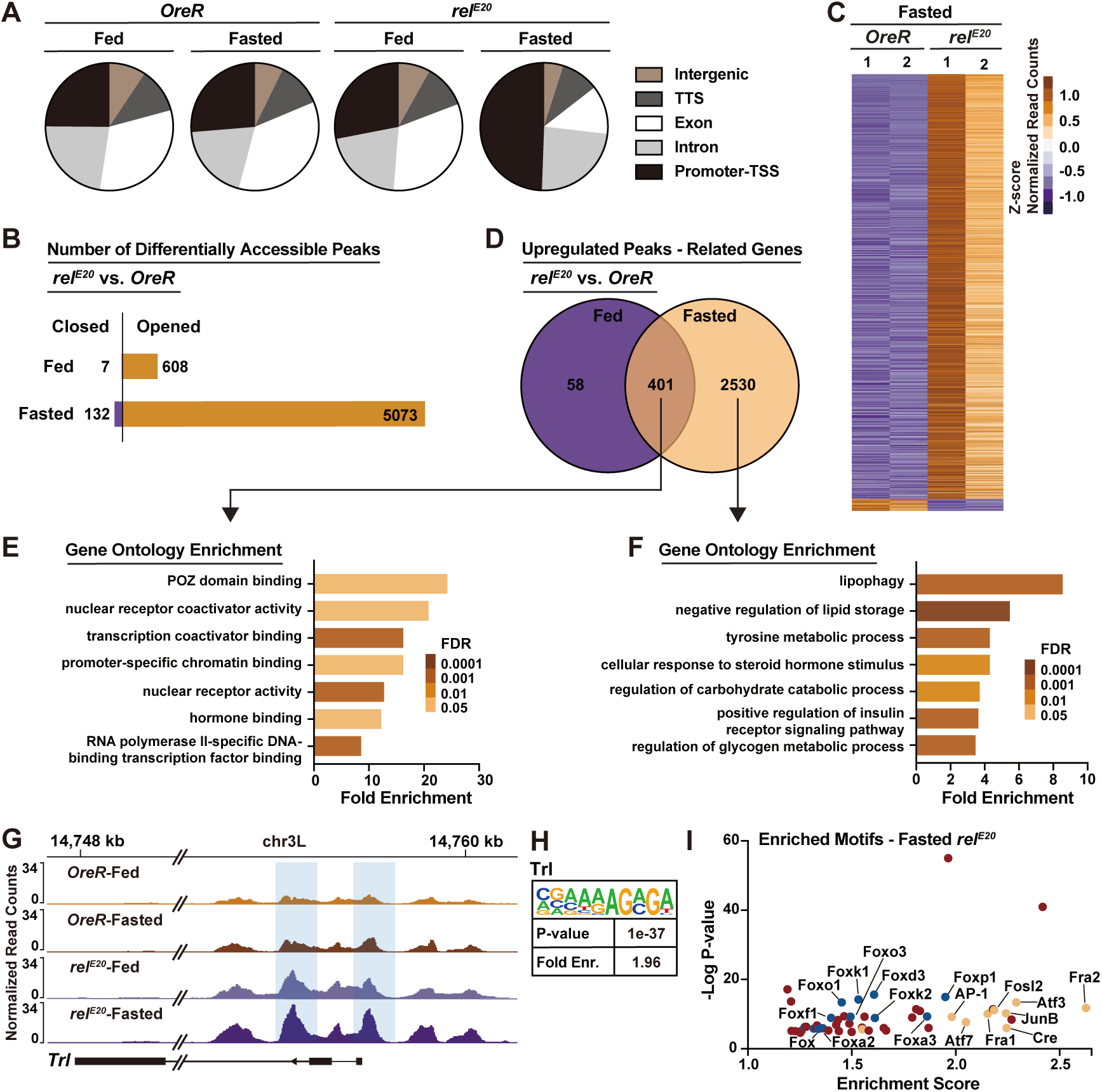
NF-κB/Relish limits fasting-induced changes in chromatin accessibility. (A) Distribution of genomic features of all accessible regions. Five annotations were examined, transcription terminal site (TTS), promoter-transcription start site (TSS), exon, intron, and intergenic. (B) The number of differentially accessible peaks in Relish mutants compared with OreR controls. (C) Heatmap of differentially accessible peaks between Relish mutants and *OreR* controls during fasting. (D) Venn diagram illustrating the overlap and distinct gene sets associated with upregulated chromatin peaks in Relish mutant flies. (E) Overlap upregulated genes analyzed for gene ontology (GO) terms. (F) Distinct upregulated genes during fasting analyzed for gene ontology (GO) terms. (G) Normalized chromatin accessibility profiles at *Trl* gene locus. Shaded regions indicate chromatin peaks exhibiting uniquely increased accessibility in Relish mutant flies. (H) Trl motif enriched in the regions gained accessibility in Relish mutants. (I) Motif enrichment analysis identifies transcription factor motifs enriched in chromatin regions that gained accessibility in fasted Relish mutant flies. The Forkhead box (Fox) family is represented by blue, and the Activator Protein-1 (AP-1) family is represented by yellow. Fasted represents 20h of nutrient deprivation. All flies were 7 days old post-eclosion.

We next compared gene sets associated with peaks that gained accessibility in *rel^E^*^20^ homozgyous mutants under *ad libitum* feeding and fasting conditions. During *ad libitum* feeding, there were 608 peaks that showed increased chromatin accessibility in NF-κB/Relish mutant flies (Figure 1B), which were associated with 459 genes (Figure 1D). However, during acute fasting the number of corresponding genes associated with differentially accessible peaks was significantly increased to 2,931 (Figure 1D). These two gene sets shared 401 overlapping genes, suggesting that they are intrinsically regulated by NF-κB and are not affected by changes in nutrient deprivation. Moreover, 2,530 genes uniquely gained accessibility during nutrient deprivation, indicating that these genes are subject to fasting-induced genomic remodeling and are normally restrained by the presence of NF-κB/Relish. Gene ontology (GO) annotation indicated that common (overlapping in *ad libitum* and fasting conditions) genes were predominately enriched in pathways related to transcription and chromatin binding (Figure 1E). For instance, the promoter-TSS region of the pioneer transcription factor Trl (Trithorax-like) gained accessibility in NF-κB/Relish mutant flies (Figure 1G). Additionally, we found a significant enrichment of Trl binding motifs in peaks that were more accessible in NF-κB/Relish mutants during *ad libitum* feeding (Figure 1H), demonstrating that Trl may contribute to the intrinsic chromatin accessibility changes caused by the loss of NF-κB function.

GO annotation also indicated that genes which gained accessibility exclusively during fasting in Relsih mutants (Figure 1D) were enriched for a wide variety of cellular processes and functions, including epithelial cell migration and morphogenesis, metabolic processes, humoral immune responses, organelle localization, RNA regulation, nucleolus regulation, and negative and positive regulation of a diverse array of signaling pathways. Using a biased analysis towards metabolic processes, we uncovered that metabolic genes which gained accessibility exclusively during fasting were mainly involved in catabolism-driven metabolic pathways (Figure 1F), such as lipolysis, lipophagy, tyrosine (amino acid) catabolism, and carbohydrate catabolism, consistent with the broad activation of metabolic transcriptional programs induced by nutrient deprivation. Thus, NF-κB/Relish appears to restrict chromatin accessibility during metabolic adaptation to nutrient changes and potentially limit the inducibility of metabolic genes associated with energy catabolism.

Previous work has suggested that NF-κB transcription factors can repress transcription through functional antagonism of other transcriptional activators^17, 18^. Our data indicates that NF-κB/Relish-dependent chromatin remodeling may be a key molecular feature of this antagonism. To uncover potential transcription factors antagonized by NF-κB/Relish during fasting, we performed Motif Enrichment analysis within open genomic regions identified in our ATAC-seq datasets. Among these regions, two well-characterized transcription factor families, Forkhead box (Fox) and Activator Protein-1 (AP-1), were highly enriched (Figure 1I). Foxo (the sole ortholog of Fox in Drosophila) is activated during starvation and is essential for the induction of catabolic gene expression^46^. This aligns with our previous finding that fasting-induced alterations in lipid metabolism are modulated through NF-κB/Relish-dependent attenuation of Foxo transcriptional activity^16^. The AP-1 transcription factor is a conserved dimeric complex typically consisting of the dFos and dJun proteins in Drosophila^47–49^. Similar to NF-κB and Foxo, AP-1 also is a rapidly inducible transcription factor^50^. While NF-κB and AP-1 are activated by distinct mechanisms, they can be activated simultaneously by the same stimuli and functionally interact to modulate transcription^51–54^.

Taken together, these findings highlight a role for NF-κB/Relish in regulating chromatin organization, particularly by restricting nutrient-dependent chromatin remodeling at metabolic target gene loci, which may be critical for metabolic adaptation.

### NF-κB/Relish restrains fasting-induced metabolic transcriptional programs through chromatin modulation

Catabolic transcriptional programs need to be tightly regulated to promote metabolic adaptation without driving dysfunction, balancing transcriptional activation and repression. Our current ATAC-seq datasets show that NF-κB/Relish broadly limits chromatin accessibility at various metabolic target genes during fasting. To determine the potential role of NF-κB/Relish in tuning nutrient-dependent metabolic gene expression, we assayed transcriptional changes of metabolic target genes identified in Figure 1F in *rel^E^*^20^ homozygous mutants. The metabolic programs shown in Figure 1F comprise a total of 36 genes, among which 18 genes were significantly repressed (restrained) by NF-κB/Relish at the whole-body level during acute fasting (Figure S2A). Notably, the triglyceride lipase *Bmm* (Brummer or Drosophila Adipose triglyceride lipase; dATGL) was among the genes repressed, supporting our previous findings^16^. These data highlight that NF-κB can limit the transcriptional inducibility of nutrient-dependent metabolic programs.

The insect fat body connects nutrient sensing and macronutrient energy expenditure, and as such is a major lipid depot that combines storage, synthesis, and breakdown functions of vertebrate adipose and hepatic tissues^55–57^. Due to its role in energy catabolism and anabolism, we further analyzed NF-κB/Relish-metabolic target gene expression (described in Figure S2A) in fat body. We identified genes that are transcriptionally restrained by NF-κB/Relish during acute fasting in fat body/adipose, including genes controlling lipolysis and lipophagy (*Atg1* [Autophagy related 1] and *Bmm)*, tyrosine metabolism/catabolism *(Tat* [Tyrosine aminotransferase], *Hpd* [4-hydroxyphenylpyruvate dioxygenase], and *Faa* [Fumarylacetoacetate hydrolase or dFAH]), as well as carbohydrate metabolism/catabolism (*Pepck2* [Phosphoenolpyruvate carboxykinase 2], *Glyp* [Glycogen phosporylase], and *Gbs-76A* [Glycogen-binding subunit 76A]), described in Figure 2A. The genomic loci of these genes also showed enhanced chromatin accessibility in NF-κB/Relish mutant animals during fasting (Figures 2B-D, and Figures S2B-C), suggesting NF-κB limits the nutrient-dependent inducibility of specific metabolic target genes by adjusting chromatin organization.

**Figure 2.**
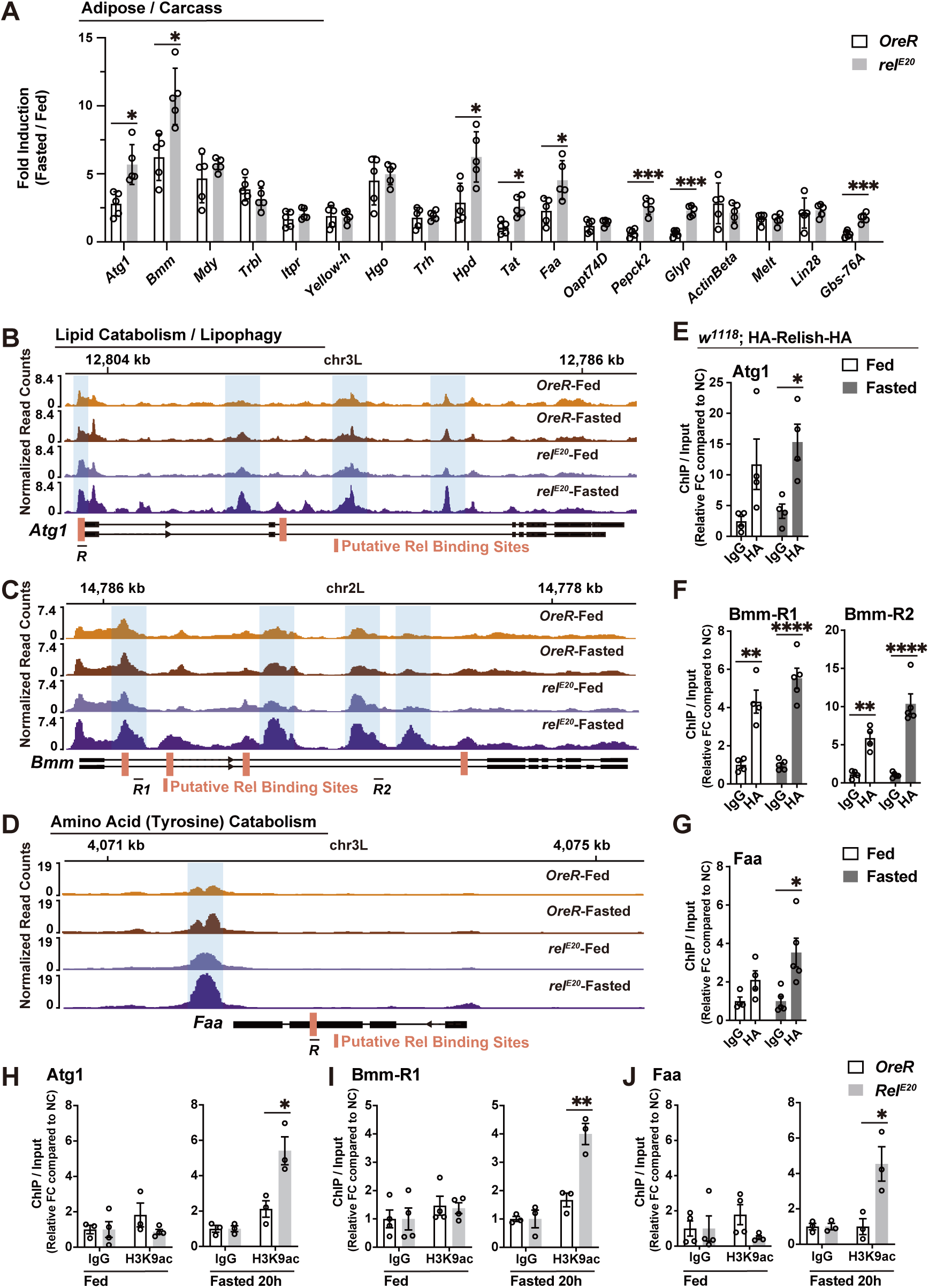
NF-κB/Relish controls fasting-induced metabolic transcriptional programs through chromatin remodeling. (A) Transcriptional changes (measured by qRT-PCR in the fat body (adipose)/carcass of OreR control and Relish mutant flies, plotted as fold induction [ratio of 20 h fasted to fed] of relative expression) of distinct upregulated metabolic genes identified in Figure S2A. n = 5 replicates. (B-D) Normalized chromatin accessibility profiles at individual targeted gene loci. Shaded regions indicate chromatin peaks exhibiting uniquely increased accessibility in fasted Relish mutant flies. Red bars represent Relish binding sites predicted by JASPAR (relative profile score threshold > 75%). *R̄* represent regional target sites (and corresponding primer sets) tested in ChIP-qPCR analysis. (E-G) ChIP-qPCR analysis of Relish binding to the predicted sites in fed and fasted (20h) conditions (genotype: *w1118*; HA-Relish-HA). ChIP-qPCR analysis with normal IgG is included as a control. Plotted as fold change (FC) of indicated PCR primer sets compared to a negative control (NC) primer set. n= 4-5 biological replicates. (H-J) ChIP-PCR analysis of H3K9ac enrichment in Relish-binding regions in wild-type (WT; OreR) and Relish mutant genotypes before and after fasting (20 h). n = 3-4 biological replicates. Bars in (A) represent mean ± SE, bars in (E-J) represent mean ± SEM. *p value < 0.05; **p value < 0.01; ***p value < 0.001; ****p value < 0.0001. All flies were 7 days old post-eclosion.

Utilizing a JASPAR database, we also identified potential NF-κB DNA binding motifs (κB sequence sites) throughout loci associated with these genes (focusing on *Atg1, Bmm, Faa, Tat,* and *Hpd*). These binding sites are located in traditional promoter and/or intronic enhancers, but also in exons, which is not uncommon in chromatin regulation, especially in animals with condensed genomes^58, 59^. To assess NF-κB/Relish DNA binding, we constructed a transgenic fly that endogenously expresses Relish containing HA tags at both the N- and C-termini (*w1118*; HA-Relish-HA). Drosophila Relish is governed by conserved upstream regulators and the IKK (IκB Kinase) signalosome (which consists of homologs of both IKKβ and IKKϒ/NEMO), while an apical caspase is required for proteolytic cleavage of the IκB domain (C-terminal), allowing for nuclear translocation and DNA binding of the active N-terminal domain^44, 60^. The proper regulation (expression and cleavage) of the HA-Relish-HA transgene in fat body was confirmed via standard infection-dependent regulation of NF-κB (Figure S3A). Utilizing the HA-epitope of this transgenic fly, we performed chromatin immunoprecipitation (ChIP)-qPCR experiments and found that NF-κB/Relish DNA binding was enriched at predicated binding motifs within these loci (*Atg1, Bmm, Faa, Tat*, and *Hpd*), particularly in fasted flies (Figures 2E-G, and Figures S3B-C). All ChIP-qPCR experiments were performed in whole flies due to inefficient chromatin extraction from dissected tissues. These NF-κB-binding motifs thus likely act as important regulatory regions to limit nutrient-dependent metabolic gene expression.

We next wanted to directly show that NF-κB/Relish can mediate epigenetic silencing of these catabolic genes through shaping histone modifications. Again using ChIP-qPCR, we monitored the level of histone 3 lysine 9 acetylation (H3K9ac, a transcriptional activation) at NF-κB/Relish regulatory regions Relish mutants (compared to OreR controls). During *ad libitum* feeding, H3K9ac enrichment was comparable between genotypes (Figures 2H-J, and Figures S3D-E). However, during nutrient deprivation, we observed elevated H3K9ac levels within regulatory regions of catabolic genes in Relish mutant flies (*Atg1, Bmm, Faa, Tat*, and *Hpd*; Figures 2H-J, and Figures S3D-E), indicating a transcriptionally active chromatin state at these genomic loci.

Altogether these data show that NF-κB/Relish can regulate a diverse array of metabolic target genes through direct DNA binding, subsequently influencing histone acetylation and chromatin remodeling at gene loci. This regulatory role enables NF-κB/Relish to repress or restrain metabolic gene transcription, limiting inducibility of nutrient-dependent gene expression linked to catabolism of energy substrates, from lipids to amino acids.

### NF-κB/Relish tunes fasting-induced catabolism to promote metabolic adaptation and enhance resilience to nutrient challenges

Metabolic adaptation to nutrient changes requires efficient allocation of macronutrient energy substrates, especially stored lipids, balancing catabolic and anabolic metabolic programs. These adaptive responses represent ancient protective mechanisms that enable metazoans across taxa to cope with food scarcity and preserve somatic stress responses, such as innate immunity. Our previous findings revealed that NF-κB/Relish can limit lipid catabolism during fasting, safeguarding the animal from excessive macronutrient energy depletion, as well as promoting appropriate metabolic adaptive responses and metabolic resilience^16^. To this end, we confirmed that *rel^E^*^20^ homozygous mutant flies exhibit increased sensitivity to fasting compared with controls (Figure 3A), accompanied by an accelerated decline in lipid storage (triglyceride; TAG) and a significant reduction in neutral lipids and/or lipid droplets in fat body during acute fasting (Figures 3B-C). We next explored roles for the nutrient-dependent metabolic/catabolic genes repressed by NF-κB/Relish, described in Figure 2, in metabolic adaptation. Bmm/dATGL is essential for lipolysis or breakdown of stored TAGs, while Atg1 contributes to lipophagy or breakdown of lipid droplets in response to nutrient deprviation^61, 62^. Tat, Hpd, and Faa are enzymes required for the breakdown of tyrosine into fumarate (a key metabolite in the TCA cycle) and acetoacetate (a ketone). Attenuation of Atg1 and Bmm specifically in fat body (CGGal4>UAS-Atg1^RNAi#1^, CGGal4>UAS-Atg1^RNAi#2^ or CGGal4>UAS-Bmm^RNAi^) enhanced survival rates and slowed lipid depletion (promoted storage) during fasting (Figures 3D and 3G). Interestingly, attenuating Tat, Hpd and Faa function in fat body (CGGal4>UAS-Hpd^RNAi#1^, CGGal4>UAS-Hpd^RNAi#2^, CGGal4>UAS-Tat^RNAi^, or PplGal4^ts^>UAS-Faa^RNAi^; a conditional genetic driver was used to attenuate Faa to avoid developmental phenotypes) also enhanced survival rates (Figures 3E-F). Attenuation of Tat and Faa also led to decelerated lipid depletion during acute fasting; Hpd attenuation, however, did not significantly affect TAG levels (Figures 3H-I). These results indicate that tyrosine catabolism, to some extent, also impinges on lipid metabolism in response to nutrient changes. Furthermore, genetically attenuating Faa in the fat body rescued the accelerated loss of lipid storage and decreased survival associated with Relish loss-of-function during fasting (PplGal4^ts^>UAS-Faa^RNAi^; *rel^E^*^20^*/rel^E^*^20^, Figures 3J-K).

**Figure 3.**
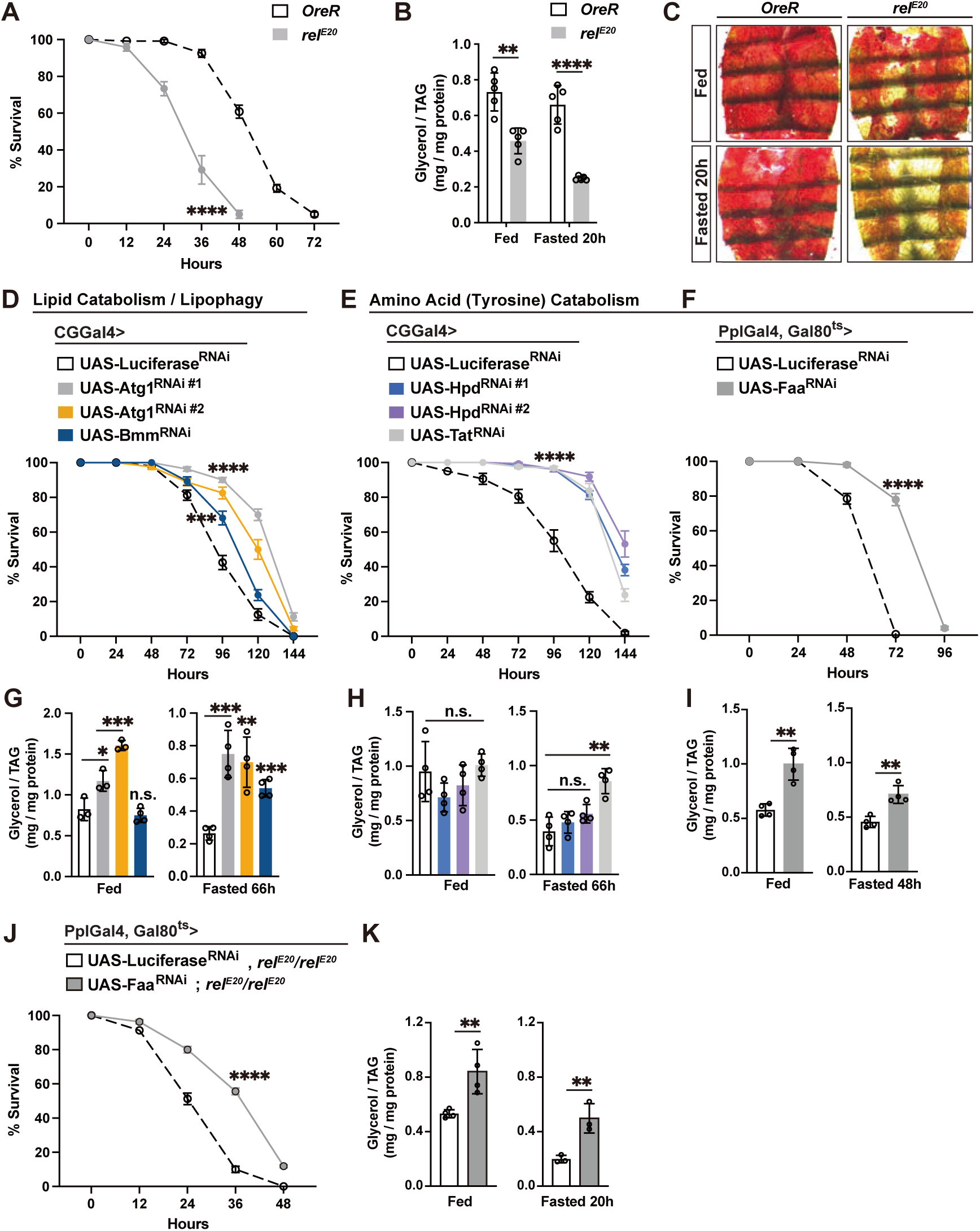
Metabolic programs adjusted by NF-κB/Relish function shape metabolic adaptation to fasting. (A-C) Relish-dependent changes in lipid metabolism and survival in response to fasting. (A) Starvation resistance of female flies. n = 6 cohorts (total 120 flies). Genotypes: OreR (wild type, WT, control) and *rel^E^*^20^*/rel^E^*^20^ (mutant). (B) Total triglyceride (TAG) levels of whole bodies before and after fasting (20 h). n = 5 samples. (C) Oil red O (ORO) stain of dissected carcass/fat body (adipose) before and after fasting (20 h). (D-I) Attenuating Relish repressed metabolic genes in fat body restores metabolic adaptation responses. (D-F) Starvation resistance of female flies (CGGal4/+; UAS-Luciferase^RNAi^, CGGal4/+; UAS-Atg1^RNAi#1^, CGGal4/+; UAS-Atg1^RNAi#2^, CGGal4/+; UAS-Bmm^RNAi^, CGGal4/+; UAS-Tat^RNAi^, CGGal4/UAS-Hpd^RNAi#1^, CGGal4/+; UAS-Hpd^RNAi#2^, PplGal4-tubGal80^ts^/+; UAS-Luciferase^RNAi^ or PplGal4-tubGal80^ts^/UAS-Faa^RNAi^). n=8-10 cohorts (total 160-200 flies). (G-I) Total TAG levels of whole bodies (n = 4 samples). (J and K) Attenuating Faa in fat body of Relish-deficient flies restores metabolic adaptation responses. (J) Starvation resistance of female flies (PplGal4-tubGal80^ts^/+; UAS-Luciferase^RNAi^, *rel^E^*^20^*/rel^E^*^20^ or PplGal4-tubGal80^ts^/UAS-Faa^RNAi^, *rel^E^*^20^*/rel^E^*^20^). n=8 cohorts (total 160 flies). (K) Total TAG levels of whole bodies (n = 3-4 samples). Bars in (A, D-F and J) represent mean ± SEM, bars in (B, G-I and K) represent mean ± SE. n.s. (no significant); *p value < 0.05; **p value < 0.01; ***p value < 0.001; ****p value < 0.0001. All flies were 7 days old post-eclosion.

Taken together, these results suggest that NF-κB/Relish mediates metabolic adaptation to nutrient changes through restraining excessive transcriptional activation of catabolic genes, tuning macronutrient energy homeostasis, and promoting metabolic resilience.

### Histone Deacetylase 6 (HDAC6) restrains fasting-induced metabolic transcriptional programs

To further investigate the mechanisms underlying NF-κB/Relish-mediated suppression of chromatin accessibility and H3K9ac histone acetylation, we performed functional genetic screens targeting protein families with histone deacetylase activity. Histone deacetylases (HDACs) can be classified into two families based on the presence of a conserved deacetylase domain and/or dependency: the classical HDAC family and Sirtuin (Sirt) protein family^63–65^. In Drosophila, five members of each family have been identified, and some are functionally linked to nutrient deprivation and/or lipid metabolism^46, 66, 67^. For example, HDAC4 and Sirt1 independently modulate hormone- and nutrient signaling-activated fasting programs by deacetylating the transcription factor Foxo^46^. Thus, to identify a potential cofactor(s) of NF-κB/Relish that could directly modulate chromatin, we performed a fat body-specific (CGGal4>UAS-Gene^RNAi^) RNAi screen targeting individual histone deacetylases and evaluated key physiological parameters (fasting survival sensitivity and lipid content) regulated by NF-κB/Relish during metabolic adaptation. Attenuating HDAC6, Sirt1 and Sirt7 in fat body (CGGal4>UAS-HDAC6^RNAi^, CGGal4>UAS-Sirt1^RNAi^ or CGGal4>UAS-Sirt7^RNAi^) resulted in increased sensitivity to fasting, however, accelerated loss of TAG was only observed in flies with attenuated HDAC6 (CGGal4>UAS-HDAC6^RNAi^, Figures 4A-D). These changes in TAG levels correlated with a significant reduction of neutral lipid storage in fat body compared to controls (Figure 4E). Conversely, attenuating HDAC4 in fat body (CGGal4>UAS-HDAC4^RNAi^) significantly limited fasting-mediated decreases in lipids and prolonged survival (Figures 4A and 4C), consistent with its role in enhancing (not limiting) Foxo-dependent lipolysis during fasting. Similar phenotypes were confirmed in HDAC6 knockout flies (*HDAC6^KO^*, Figures 4F and 4G) or using an independent fat body driver (PplGal4>UAS-HDAC6^RNAi^, Figures S4A-C). These metabolic responses in HDAC6 loss-of-function flies phenocopy metabolic responses in NF-κB/Relish loss-of-function flies, suggesting that HDAC6 may function as a cofactor in NF-κB/Relish-mediated chromatin modulation and metabolic target gene repression during nutrient deprivation.

**Figure 4.**
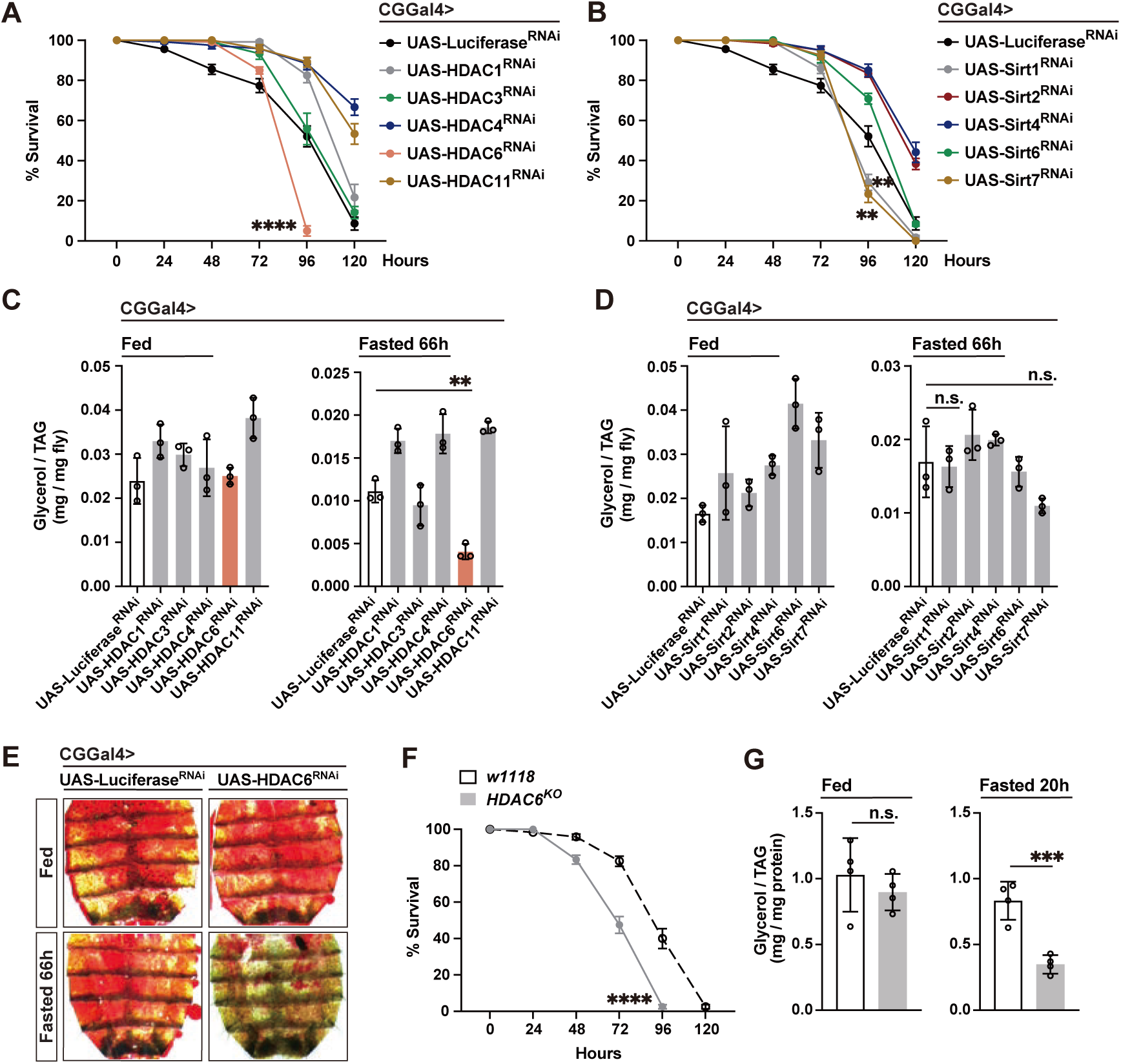
Histone deacetylase 6 (HDAC6) function directs nutrient-dependent metabolic adaptation. (A-D) Biased functional genetic screen identifies key Relish co-factor with histone deacetylase activity during starvation. (A and B) Starvation resistance of female flies (CGGal4/+; UAS-Luciferase^RNAi^, CGGal4/+; UAS-HDAC1^RNAi^, CGGal4/+; UAS-HDAC3^RNAi^, CGGal4/+; UAS-HDAC4^RNAi^, CGGal4/+; UAS-HDAC6^RNAi^, CGGal4/+; UAS-HDAC11^RNAi^, CGGal4/+; UAS-Sirt1^RNAi^, CGGal4/+; UAS-Sirt2^RNAi^, CGGal4/+; UAS-Sirt4^RNAi^, CGGal4/+; UAS-Sirt6^RNAi^ or CGGal4/+; UAS-Sirt7^RNAi^). n=6 cohorts (total 120 flies). (C and D) Total TAG levels of whole bodies (n = 3 samples). (E) ORO stain of dissected carcass/fat body (adipose) before and after fasting (66 h). (F) Starvation resistance of female flies (*w1118 [w^−^]* controls and *y^−^*,*w^−^*; *HDAC6^KO^* homozygote mutants). n=6 cohorts (total 120 flies). (G) Total TAG levels of whole bodies (n = 4 samples). Bars in (A, B and F) represent mean ± SEM, bars in (C, D and G) represent mean ± SE. n.s. (no significant); **p value < 0.01; ***p value < 0.001; ****p value < 0.0001. All flies were 7 days old post-eclosion.

HDAC6, a class IIb histone deacetylase, is well characterized as a cytosolic Zn^2+^-dependent deacetylase mainly involved in deacetylation of non-histone proteins^68–70^. In Drosophila, HDAC6 serves as a key integrator of the cytoskeletal dynamics and proteostasis systems in the cytoplasm^71–73^. However, it has an emerging role in the nucleus to shape chromatin, especially in vertebrates^74, 75^. To elucidate the function of HDAC6 in regulating nutrient-dependent metabolic transcriptional programs, we first showed that we could rescue the accelerated lipid depletion and increased sensitivity to fasting found in HDAC6-attenuated flies by re-expressing HDAC6 in fat body (CGGal4>UAS-HDAC6^RNAi^/UAS-HDAC6-HA; Figures 5A-C). This dysfunctional metabolic adaptation can also be rescued by re-expressing human HDAC6 in fat body (PplGal4>UAS-hHDAC6; UAS-HDAC6 ^RNAi^, Figures S4D-F), which has been shown to have key nuclear functions. We also uncovered that, like NF-κB/Relish (Figure 2A), HDAC6 is required to restrain excessive transcriptional activation (or limit inducibility) of catabolic genes during acute fasting. *Atg1, Bmm, Faa, Tat*, *Hpd, Pepck2, and Glyp* transcription was significantly upregulated in fat body-specific HDAC6-attenuated flies during acute fasting and could be restored by re-expressing HDAC6 in fat body/adipose (CGGal4>UAS-HDAC6^RNAi^/UAS-HDAC6; Figures 5D-G). Only *Gbs-76A* transcription exhibited no change across these genetic backgrounds. Finally, as a proof-of-concept, genetically restricting lipid catabolism in fat body by targeting Bmm/dATGL (CGGal4>UAS-HDAC6^RNAi^/UAS-Bmm^RNAi^) can rescue maladaptive metabolic responses caused by attenuated HDAC6 function and promote metabolic resilience (Figures S4G-I).

**Figure 5.**
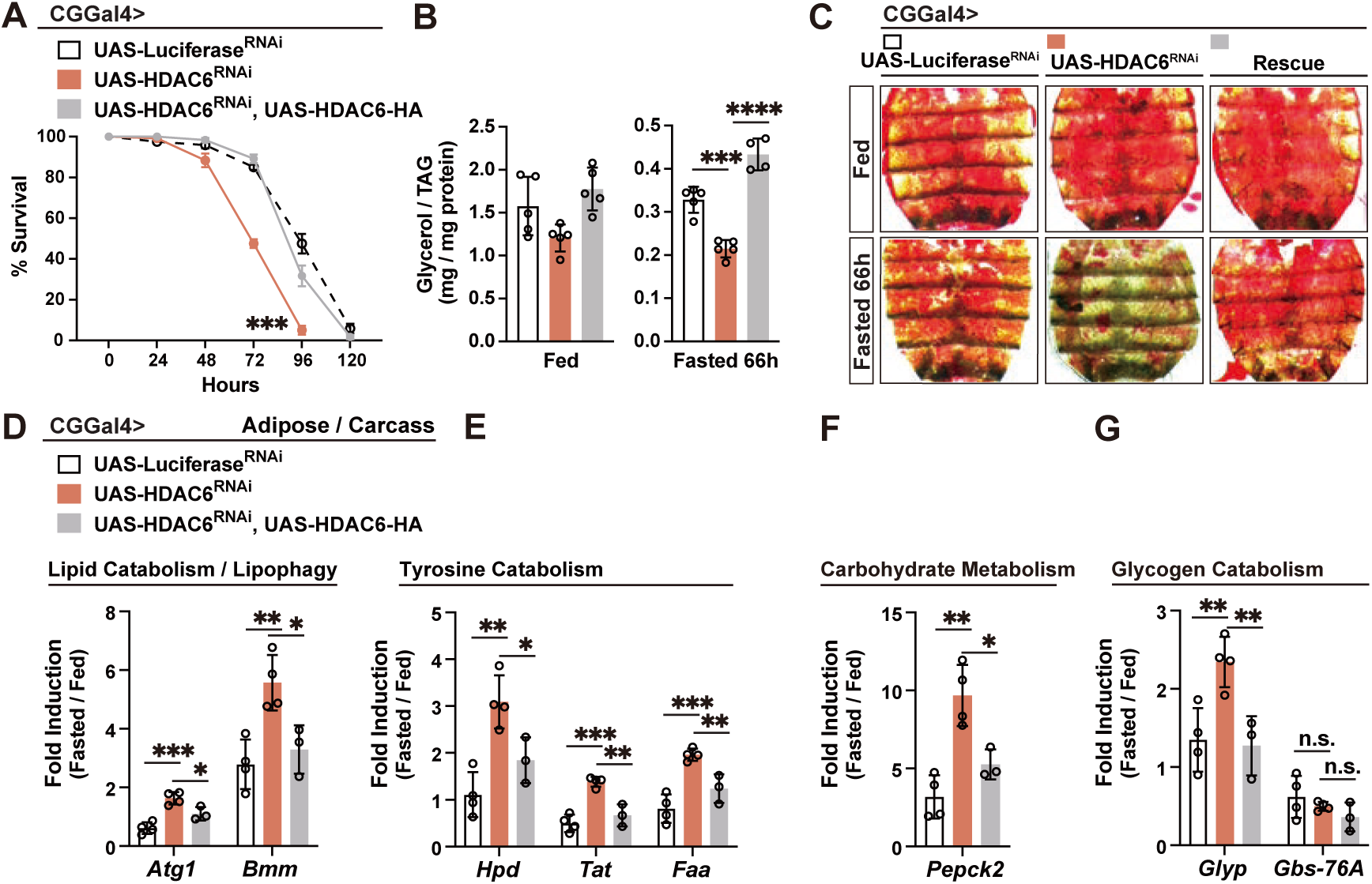
HDAC6 regulates fasting-induced metabolic transcriptional programs. (A-C) Re-expressing HDAC6 in the fat body of HDAC6-deficient flies restores metabolic adaptation responses. (A) Starvation resistance of female flies (CGGal4/+; UAS-Luciferase^RNAi^, CGGal4/+; UAS-HDAC6^RNAi^ or CGGal4/+; UAS-HDAC6^RNAi^/UAS-HDAC6). n=6 cohorts (total 120 flies). (B) Total TAG levels of whole bodies (n = 4-5 samples). (C) ORO stain of dissected carcass/fat body (adipose) before and after fasting (66 h). (D-G) Transcriptional changes (measured by qRT-PCR, plotted as fold induction [ratio of 20 h fasted to fed] of relative expression) of metabolic genes repressed by Relish in the fat body (adipose)/carcass. n = 3-4 replicates. Bars in (A) represent mean ± SEM, bars in (B, D-G) represent mean ± SE. n.s. (no significant); *p value < 0.05; **p value < 0.01; ***p value < 0.001; ****p value < 0.0001. All flies were 7 days old post-eclosion.

These data show that HDAC6, like NF-κB/Relish, is required to limit the inducibility of nutrient-dependent metabolic transcriptional programs, restrain catabolic responses, and tune metabolic adaptation. HDAC6 may thus function as a crucial cofactor in NF-κB/Relish-mediated suppression of catabolic gene expression, histone acetylation, and chromatin remodeling.

### HDAC6 interacts with NF-κB/Relish to shape nutrient-dependent chromatin remodeling

We next investigated the relationship between HDAC6 and NF-κB/Relish in dictating nutrient-dependent epigenetic regulation of catabolic target genes, first exploring a role for nuclear HDAC6. Using a transgenic fly endogenously expressing sGFP- and Flag-tagged HDAC6 (referred to as *w1118*; HDAC6-sGFP-TVPTBFlag [FlyFos]), we fractionated the cytoplasmic and nuclear components of dissected fat body/adipose and found that HDAC6 is predominantly localized in the cytoplasm, but with a minor fraction present in the nucleus (Figure 6A, similar to ratios found vertebrate cells)^76^. In contrast, NF-κB/Relish was distributed evenly between the cytoplasm and nucleus. Notably, the subcellular distribution of both proteins remained unchanged under different nutritional conditions. Thus, Relish-dependent repression (as opposed to canonical transcriptional activation) of gene expression may represent a constitutive or steady-state function of NF-κB (described in vertebrates ^77–79^), and thus could act as a priming mechanism that promotes metabolic resilience in response to nutrient stress. However, the nuclear localization of HDAC6 implies a potential role in the regulation of nutrient-dependent chromatin changes.

**Figure 6.**
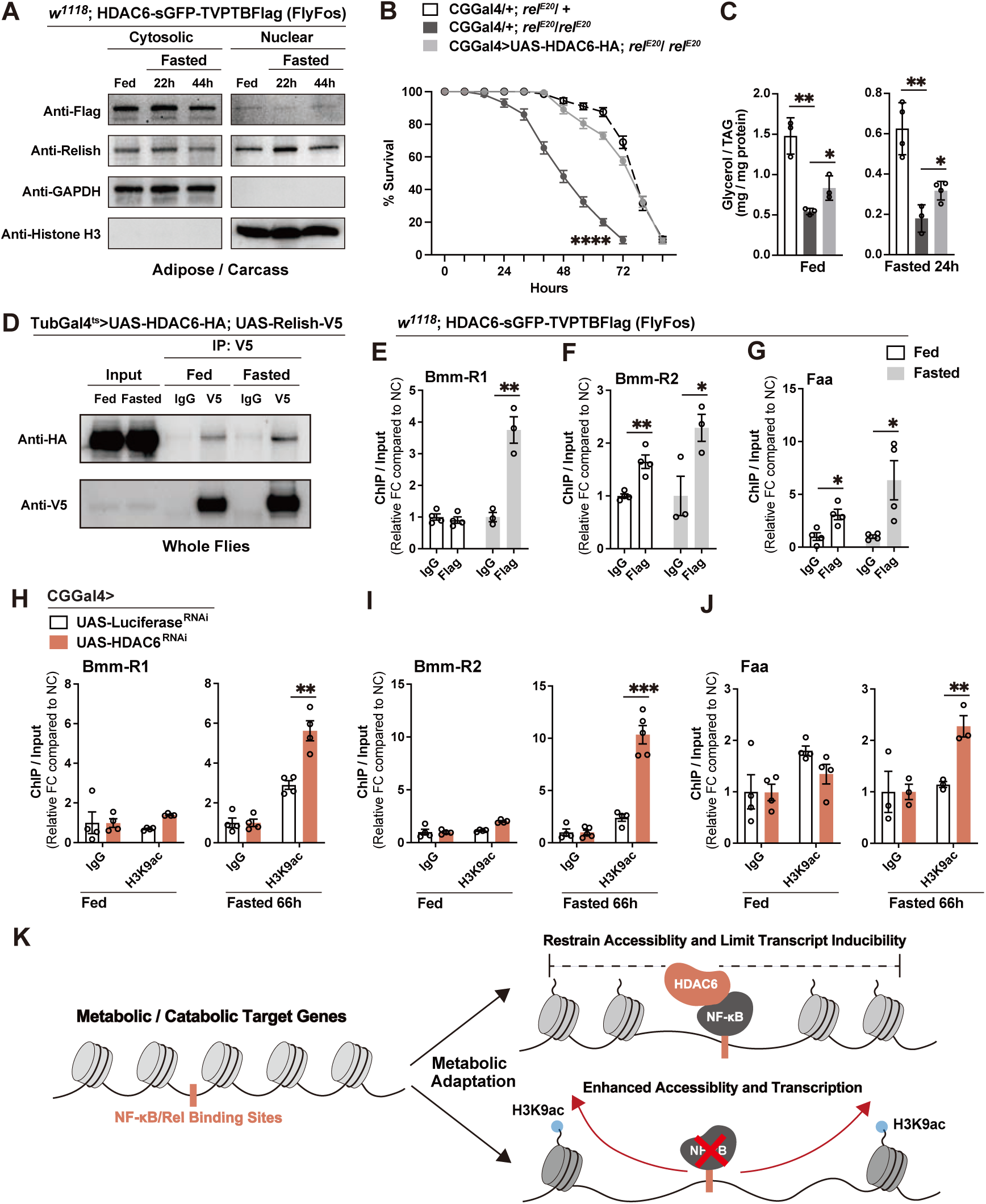
Genetic and physical interactions between NF-κB/Relish and HDAC6 direct nutrient-dependent changes in chromatin accessibility. (A) The protein levels of Relish and HDAC6 in the cytosol and nuclear of adipose (fat body) before and after starvation. GAPDH and Histone H3 were used as the markers of cytoplasm and nuclear, respectively. Genotype: *w1118*; HDAC6-sGFP-TVPTBFlag. (B and C) Re-expressing HDAC6-HA in the fat body of Relish-deficient flies partially restores metabolic adaptation responses. (B) Starvation resistance of female flies (CGGal4/+; *rel^E^*^20^/+, CGGal4/+; *rel^E^*^20^*/rel^E^*^20^ or CGGal4/UAS-HDAC6-HA; *rel^E^*^20^*/rel^E^*^20^). n=10 cohorts (total 200 flies). (C) Total TAG levels of whole bodies (n = 3-4 samples). (D) *In vivo* interaction of Relish with HDAC6. Immunoprecipitates were prepared from lysates of whole flies from TubGal80^ts^/UAS-HDAC6-HA; TubGal4/UAS-Relish-V5. (E-G) ChIP-qPCR analysis of HDAC6 binding to the Relish-binding sites in fed and fasted (20h) conditions (genotype: *w1118*; HDAC6-sGFP-TVPTBFlag). n= 3-4 biological replicates. (H-J) ChIP-PCR analysis of H3K9ac enrichment in Relish-binding regions in CGGal4/+; UAS-Luciferase^RNAi^ and CGGal4/+; UAS-HDAC6^RNAi^ genotypes before and after fasting (66 h). n = 3-5 biological replicates. (K) Model summarizing conclusions. Bars in (B, E-J) represent mean ± SEM, bars in (C) represent mean ± SE. *p value < 0.05; **p value < 0.01; ***p value < 0.001; ****p value < 0.0001. All flies were 7 days old post-eclosion.

We next explored potential genetic and physical interactions between HDAC6 and NF-κB/Relish. First, we employed an *in vitro* approach and transiently expressed HA-tagged HDAC6 (pAC5.1-HDAC6-HA) and V5-tagged Relish (pAC5.1-Relish-V5) in Drosophila S2R+ cells. HDAC6 co-immunoprecipitated with Relish under serum starvation conditions, but not under normal culture conditions (Figure S5A). These results prompted us to investigate this physical interaction *in vivo*. To this end, we generated two UAS-dependent transgenic flies overexpressing C-terminally tagged HDAC6 (UAS-HDAC6-HA) and NF-κB/Relish (UAS-Relish-V5), respectively. Overexpression of HDAC6-HA in fat body (CGGal4>UAS-HDAC6-HA; *rel^E^*^20^*/ rel^E^*^20^) can partially rescue the accelerated loss of lipid storage and reduced survival in NF-κB/Relish mutant flies during fasting (Figures 6B-C). These findings suggest that the presence of NF-κB/Relish is required for HDAC6 to fully exert its role in promoting metabolic adaptation. Additionally, co-immunoprecipitation from whole-fly lysates (TubGal4^ts^>UAS-HDAC6-HA; UAS-Relish-V5) confirmed that HDAC6 physically associates with Relish *in vivo* during both *ad libitum* feeding and fasting conditions (Figure 6D). These genetic and physical interactions establish a direct molecular link between HDAC6 and NF-κB/Relish in coordinating nutrient-dependent metabolic adaptative responses.

The nuclear localization of HDAC6, and its interaction with NF-κB/Relish, led us to further explore HDAC6’s regulatory role at NF-κB/Relish DNA binding sites in catabolic target gene loci (Figure 2). To precisely assay DNA interactions, we used *w1118*; HDAC6-sGFP-TVPTBFlag transgenic flies to avoid potential nuclear accumulation caused by HDAC6 overexpression. Utilizing ChIP-qPCR, we found that HDAC6 was significantly enriched at NF-κB/Relish binding motifs within metabolic target gene loci (*Bmm, Tat, Faa,* and *Hpd* were tested), especially during acute fasting (Figures 6E-G and S5B-C). We next determined if HDAC6 could influence histone acetylation and chromatin remodeling within these catabolic genes. While HDAC6-DNA interactions were detected under *ad libitum* feeding conditions in many cases (Figures 6E-G and S5B-C), attenuating HDAC6 in fat body (CGGal4>UAS-HDAC6^RNAi^) did not alter the H3K9ac levels at these loci (employing ChIP-qPCR; Figures 6H-J and S5D-E). During fasting, however, fat body-specific attenuation of HDAC6 (CGGal4>UAS-HDAC6^RNAi^) resulted in a significant enrichment (compared to controls) of H3K9ac levels at NF-κB/Relish regulatory motifs within these genes (Figures 6H-J and S5D-E, similar to *rel^E^*^20^ mutants), also suggesting that HDAC6- and NF-κB/Relish-mediated chromatin remodeling is fasting-dependent. These findings suggest that HDAC6 acts as a cofactor of NF-κB/Relish to restrain nutrient-dependent metabolic transcriptional programs through dampening chromatin accessibility and thus balancing catabolic responses.

## Discussion

In summary, we uncovered a distinct mechanism by which the NF-κB transcription factor Relish in Drosophila fine-tunes metabolic adaptation through chromatin regulation, restraining excessive metabolic transcriptional activation in response to nutrient changes (Figure 6K). Our findings show that NF-κB transcription factors, classically known for their roles in innate immunity, are also critical guardians of macronutrient energy balance. The absence of NF-κB/Relish leads to exacerbated catabolic responses to nutrient deprivation, from enhanced and widespread chromatin accessibility to heightened metabolic gene expression, resulting in rapid depletion of macronutrient energy storage and reduced resilience. By partnering with HDAC6 to modulate histone acetylation and restricting open chromatin, NF-κB/Relish limits the inducibility of metabolic target genes when nutrients are scarce. These findings highlight NF-κB transcription factors as key integrators of immune and metabolic signaling architecture, promoting balanced adaptive metabolic responses.

Balancing metabolic responses requires a symmetry between conserving macronutrient energy storage with usage, as well as promoting other somatic stress responses. Nutrient deprivation forces metazoans to reprioritize macronutrient energy usage, often at the expense of innate immunity. For example, acute starvation increases susceptibility to bacterial infection in Drosophila^80^. This trade-off likely reflects, in part, the high energy demand of immune responses. Proper regulation of macronutrient energy mobilization during starvation is therefore critical. An overzealous catabolic response can deplete vital reserves, whereas insufficient mobilization impairs acute survival. We show that Relish (a Drosophila NF-κB homolog) acts as a rate-limiting brake on fasting-induced energy mobilization by repressing transcriptional overactivation of metabolic genes. It’s thus possible that NF-κB transcription factors temper the conflict between ‘survival’ metabolism and immunity by preventing “too much, too fast” consumption of stored energy during nutrient challenges. This concept dovetails with emerging recognition that NF-κB pathways link metabolism and immunity across taxa^9^. In mammals, for instance, NF-κB signaling in immune cells actively reprograms cellular metabolism during inflammation and dysregulation of this crosstalk can lead to metabolic diseases^81^. Our study extends this paradigm, suggesting that proper energy balance during nutrient stress may, in turn, feedback to influence immune competence, highlighting the evolutionary conservation of immune-metabolic coupling.

This study also revealed a genome-wide surge in chromatin accessibility in Drosophila Relish mutants during acute fasting. Over 5,000 chromatin regions (associated with ∼2,500 genes) become accessible upon starvation in the absence of Relish, compared to only ∼130 regions in wild-type flies (Figures 1B and 1D), indicating that NF-κB/Relish dynamically shapes the chromatin landscape during nutrient stress. This suggests the NF-κB proteins are not only sequence-specific transcription factors, but also chromatin modulators, influencing which genomic regions are accessible for transcription upon fasting. Such a role is concurrent with growing evidence that NF-κB family members globally orchestrate chromatin remodeling during stress responses. In mammalian macrophages, recent studies have highlighted that NF-κB subunits Rel-A and c-Rel are broadly required for inducible nucleosome remodeling at hundreds of latent enhancers upon Toll-like receptor stimulation^36^. Similarly, activated NF-κB prompts rapid formation of super enhancers in endothelial cells during proinflammatory stimulation^82^. Importantly, our current work extends this chromatin modulatory role for NF-κB transcription factors to the context of metabolic adaptation and resilience, thereby highlighting a fundamental mechanism of immune-metabolic regulatory crosstalk.

The ability of Drosophila NF-κB/Relish to modulate nutrient-dependent chromatin accessibility is likely achieved through interactions with the histone deacetylase HDAC6. Unlike classical nuclear HDACs, HDAC6 predominantly localizes to the cytoplasm under basal conditions. However, emerging evidence has revealed that HDAC6 can shuttle into the nucleus and influence chromatin^74, 83^. Pharmacological inhibition of HDAC6 in cancer cells increased global acetylation of histone H3 at lysines 9, 14 and 27 and conferred widespread chromatin accessibility changes^75^. Consistently, we observed that Drosophila HDAC6 is also localized in the nucleus, where it regulates histone modifications and gene expression during metabolic adaptation to fasting. Our data highlight HDAC6 as a novel metabolic stress-responsive deacetylase that, in contrast to HDAC4 and Sirt1^46^, functions to buffer excessive transcriptional activation. Under fasting conditions, the functional heterogeneity of deacetylases underscores the critical importance of maintaining energy balance and promoting metabolic resilience, which requires a precise, hierarchical, and coordinated regulatory strategy involving multiple deacetylases. To this end, our results also suggest that NF-κB-HDAC interactions impact cooperative and competitive interactions among transcription factors. Crosstalk between NF-κB and heterologous transcription factors through direct binding or occupancy at adjacent sites profoundly influences transcriptional responses^84^, which includes stress-responsive transcription factors that drive macronutrient energy catabolism, such as Foxo and/or AP-1 (Jun/Fos) transcription^16, 36, 85–88^. NF-κB-HDAC6 interactions that shift the chromatin landscape may provide a plausible mechanism for the functional antagonism between NF-κB transcription factors and other transcriptional activators that shape metabolic adaptation.

## Materials and Methods

### Drosophila husbandry and strains

The following strains were obtained from the Bloomington Drosophila Stock Center: *OreR* (Oregon-R-C, #5), *rel^E^*^20^*/rel^E^*^20^ (#9457), *w1118* (#3605), *HDAC6^KO^* (#51182), TubGal4 (#5138), Tubulin (tub)-Gal80 (temperature sensitive)-ts (#65406), PplGal4 (#58768), UAS-Luciferase^RNAi^ (#31603), UAS-Atg1^RNAi #1^ (#26731), UAS-Atg1^RNAi #2^ (#35177), UAS-Bmm^RNAi^ (#25926), UAS-Tat^RNAi^ (#76065), UAS-Hpd ^RNAi #1^ (#52923), UAS-Hpd ^RNAi #2^ (#80421), UAS-Faa^RNAi^ (#51796), UAS-HDAC1^RNAi^ (#31616), UAS-HDAC3^RNAi^ (#31633), UAS-HDAC4^RNAi^ (#28549), UAS-HDAC6^RNAi^ (#31053), UAS-HDAC11^RNAi^ (#32480), UAS-Sirt1^RNAi^ (#31636), UAS-Sirt2^RNAi^ (#31613), UAS-Sirt4^RNAi^ (#31638), UAS-Sirt6^RNAi^ (#31399), UAS-Sirt7^RNAi^ (#31093), UAS-hHDAC6 (#76817). *w1118*; HDAC6-sGFP-TVPTBFlag [FlyFos] (#318243) was obtained from Vienna Drosophila RNAi Center. CGGal4 was kindly provided by C. Thummel.

*w1118*; HA-Relish-HA, *w1118*; UAS-HDAC6-HA, *w1118*;; UAS-Relish-V5 transgenic flies were generated for this study. The ebony mutation/marker *e^S^* was removed from the *rel^E^*^20^ mutant background, with the *rel^E^*^20^ mutation finally outcrossed into a wild-type (*OreR*) background. Transgenic lines were backcrossed 10 times into the *w1118* background that was used as a control strain, with continued backcrossing every 6-8 months to maintain isogeneity.

All flies were reared on a standard yeast and cornmeal-based diet at 25 °C and 65% humidity on a 12 hr light and dark cycle, unless otherwise indicated. The standard lab diet (cornmeal-based) was made with the following protocol: 14g agar, 165.4g malt extract, 41.4g dry yeast, 78.2g cornmeal, 4.7ml propionic acid, 3g methyl 4-hydroxybenzoate and 1.5L water. To standardize metabolic results, 50 virgins were crossed to 10 males and kept in bottles for 2-3 days to lay enough eggs. Progeny of crosses were collected for 3-4 days after initial eclosion. Collected progeny (males and females) were then transferred to new bottles and allowed to mate for 2 days (representing unique populations fed *ad libitum*). Next, post-mated females (20 flies per vial/cohort) were sorted into individual vials for 2 days.

All experiments presented in the results were done utilizing female flies 7 days old post-eclosion (following dietary protocol referenced above).

### Generation of transgenic flies

*w1118*; HA-Relish-HA transgenic flies were generated by PCR amplification of the Relish sequence from *w1118* genomic DNA, with specific primers (upstream-F: TTGCTCTGGGGCTTCATCGC upstream-R: AACATCGTATGGGTACATAGTCCGTATCAATCCAATATTTCTG, gene body-F: TACCCATACGATGTTCCAGATTACGCTAACATGAATCAGTACTACGACCTG, gene body-R: AGCGTAATCTGGAACATCGTATGGGTAAGTTGGGTTAACCAGTAGGG, downstream-F: GTTCCAGATTACGCTTGATTATGTATTGAATGTTGATCAATAA, downstream-R: GCGCATAAGTACATGTATGAT), and then cloned into a pBID plasmid. This plasmid was injected into *w1118* embryos (Rainbow Transgenic Flies).

*w1118*; UAS-HDAC6-HA transgenic reporter flies were generated by PCR amplification of HDAC6 coding sequence with specific primers (F: AACAGATCTGCGGCCGCGGCTCGAGGGTACATGAGCCCGCCCATCGTC ACGC, R: GGTTCCTTCACAAAGATCCTCTAGAGTCAAGCGTAATCTGGAACATCGTA) from UGO01255 (DGRC Stock 1663924; https://dgrc.bio.indiana.edu//stock/1663924; RRID: DGRC_1663924), and then cloned into a pUASt plasmid. This plasmid was injected into *w1118*; attp40 embryos (Rainbow Transgenic Flies).

*w1118*; UAS-Relish-V5 transgenic reporter flies were generated by PCR amplification of Relish coding sequence with specific primers (F: AAGAGAACTCTGAATAGGGAATTGGGAATTCGACTACAAGGATGACGAT GACAAA, R: GGTTCCTTCACAAAGATCCTCTAGAGGTACCTCAATGGTGATGGTGATGA TGACC) from pMT-Flag-RELISH (Addgene plasmid # 63750; http://n2t.net/addgene:63750; RRID: Addgene_63750), and then cloned into a pUASt plasmid. This plasmid was injected into *w1118*; attp40 embryos (Rainbow Transgenic Flies).

### Conditional expression of UAS-linked transgenes

The TARGET system was used in combination with PplGal4 or TubGal4 to conditionally express UAS-linked transgenes in the fat body (PplGal4, tub-Gal80^ts^) or whole body (tub-Gal80^ts^; TubGal4). Flies were developed at 18 °C then shifted to 29 °C for 7 days to induce transgene expression post-eclosion.

### Generation of ATAC-seq libraries

ATAC-seq libraries were prepared as described previously^89^. In brief, approximately 120 adult female flies’ whole bodies (5-10 days old post-eclosion) were ground in liquid nitrogen then homogenized and lysed in cold lysis buffer (containing 10 mM Tris-HCl, pH 7.4, 10 mM NaCl, 3mM MgCl2 and 0.1% IGEPAL CA-630) to obtain the nuclei for each sample. Later, the Nextera DNA Sample Preparation Kit (Illumina) was adopted for carrying out the Tn-5 transposition reaction at 37 C° for 30 min. Then, DNA fragments were purified through the MinElute Kit (Qiagen) and amplified for 5 cycles using those previously designed primers containing both compatible adaptors and barcodes. Subsequently, the resulting ATAC-seq libraries were purified (MinElute Kit, Qiagen), and the final libraries were sequenced on the Illumina HiSeq4000 platform with a paired-end read of 150 bp. Raw and/or analyzed datasets are available upon request.

### Primary processing of ATAC-seq data

ATAC-seq data were processed using the standard ATAC-seq pipeline (including quality control, trimming, filtering, aligning and peaks calling) after mild modifications. In brief, the raw sequencing reads were initially processed by the FastQC program for quality control, and later the sequencing adaptors and poor-quality reads were removed using Trim Galore. Afterwards, the filtered reads were mapped to the reference Drosophila genome using BWA. Then, the sam files were converted into bam format using Samtools. To refine sequencing data, several filtering steps are implemented: (1) Reads originating from mitochondrial DNA are excluded to ensure a more accurate duplication rate assessment and to eliminate potential bias in subsequent analyses; (2)

Picard’s MarkDuplicates tool is employed to remove duplicate reads, addressing overrepresentation caused by PCR amplification; (3) Reads aligned to known blacklisted genomic regions are discarded to prevent the inclusion of problematic or non-informative data in the analysis. MACS3 was adopted for peaks calling, and an initial threshold q-value of 0.01 was used as the cutoff value. Epic2 was used to identify broad genomic domains. Utilizing deepTools, we convert filtered bam files into bigwig format, employing Counts Per Million (CPM) as the default normalization method to prepare the data for such comparative analyses and further bioinformatics processing.

### Analysis of differential accessible peaks

To generate a consensus set of unique peaks, two replicate ATAC-seq peaks were merged according to the distance between proximal end of < 1bp. Then, the average score of bigwig was further calculated across each peak using the UCSC tool. DESeq2 was employed to identify those differential accessible peaks between two different groups, with the threshold log2 fold change (FC) of > 1 and adjust p-value of < 0.05.

### Analysis of gene expression

Total RNA from the whole bodies or dissected fat body/carcass (with all of the eggs and intestines removed) of flies were extracted using Trizol and complementary DNA were synthesized using Superscript III (Invitrogen). Quantitative Real-time PCR (qRT-PCR) was performed using SYBR Green, the Thermo Fisher QuantStudio 5 Real Time PCR system, and the primers pairs described in Table S1. Results are average ± standard error of at least three independent biological samples, and quantification of gene expression levels calculated using the △Ct method and normalized to *actin5C* expression levels.

### Chromatin immunoprecipitation (ChIP) analysis

Approximately 120 adult female flies (5-10 days old post-eclosion) were ground in liquid nitrogen then homogenized and cross-linked (10 minutes at RT) in 1 mL of 1xPBS containing 1% formaldehyde, 1mM PMSF and 1x Protease Inhibitor cocktail (Thermo Scientific). The homogenate was centrifuged for 20 min at 12,000 x rpm (4 °C). The pellet was washed twice by resuspending in 1 mL of 1x PBS containing 1mM PMSF and 1x Protease Inhibitor cocktail and centrifuged at 12,000 x rpm for 20 min (4 °C). To lyse tissue and cells, the pellet was resuspended in 1,200 μL of RIPA buffer (10 mM Tris-HCl, pH 7.6, 1 mM EDTA, 0.1% SDS, 0.1% Na-Deoxycholate, 1% Triton X-100, 1mM PMSF and 1x Protease Inhibitor cocktail) then incubated at 4 °C for 30 min.

The chromatin was sheared to 500-1,000 bp DNA fragments using a Diagenode sonicator (20 min sonication, highest power, 30 sec sonication, 30 sec rest). After sonication, the sheared chromatin was centrifuged for 20 min at 12,000x rpm, 4 °C. The supernatant was collected, aliquoted, snap-frozen, and stored at -80 °C.

For immunoprecipitation, 40 μL of ChIP-grade Protein A/G Magnetic Beads (Thermo Scientific) were conjugated (4 hours incubation at 4 °C) with 10 μL of Rabbit anti-HA (Cell signaling technology, 3724S), or 2 μL of Rabbit anti-Histone H3 (acetyl 9) antibody (abcam, ab4441), or 5 μL of Mouse anti-Flag antibody (Sigma Aldrich, F1804), or 10 μL of Rabbit Control IgG (Abclonal, AC005), or 5 μL of Mouse Control IgG (Abclonal AC011). After applying beads to the magnet and removing supernatant, 1 mL of chromatin was incubated overnight with beads. Beads were washed with following buffers at 4 °C, for 10 min each: 2x with 1 mL of RIPA buffer + 1 mM PMSF + 1x protease inhibitor; 2x with 1 mL of RIPA buffer + 0.3 M NaCl; 2x with 1 mL of LiCl buffer (0.25 M LiCl, 0.5% Triton X-100, 0.5% NADOC); 1x with 1 mL of 1x TE + 0.2 Triton X-100; 1x with 1 mL of 1x TE.

To reverse crosslink, beads were re-suspended in 100 μL of 1x TE + 3 μL 10% SDS + 5 μL of 20mg/mL Proteinase K (VWR) and incubated at 65 °C overnight. Beads were applied to the magnet and supernatant was transferred to a PCR purification column (Qiagen PCR purification kit) to purify DNA. To prepare Input (chromatin extract without Immunoprecipitation), 50 μL of chromatin extract were incubated with proteinase K then applied to PCR purification column. For all Immunoprecipitated (IP) and Input samples, DNA was eluted in 30 μL of water, and 1 μL of that was used as template for qRT-PCR (see Table S1 for primer sets). Mouse or Rabbit Control IgG were used as controls.

To assess enrichment, %Input was calculated first (between ChIP DNA and input DNA for each primer set). Then the fold change in enrichment was calculated by dividing %Input of each primer set to %Input of a negative control primer set designed for Drosophila (Drosophila Negative Control primer set 1, Active Motif).

### Triglyceride (TAG) measurements in adult flies

For TAG assays, five flies (without head, per sample) were homogenized in 200 μL of PBST (PBS, 0.1% Tween 20) and heated at 70 °C for 5 min to inactivate endogenous enzymes. Samples were centrifuged at 4000 rpm for 3 min and 10 μL of cleared extract was used to measure triglycerides (StanBio Liquicolor Triglycerides Kit) according to the manufacturer instructions. TAG levels were normalized to weight or protein levels. Note: The kit measures glycerol cleaved from TAG and diacylglycerol, as well as minimal amounts of free glycerol; the majority of neutral lipids extracted from whole flies are TAG.

### Starvation sensitivity analysis

Adult flies (20 flies per vial/cohort) were provided with only water (absolutely no food) on filter paper with a KimWipe, ensuring water was present throughout the analysis. Flies were flipped into new vials every 2 days. The number of dead flies in each vial was recorded, and data is presented as the mean survival of cohorts.

### Oil Red O staining

Fat body/carcasses (with all of the eggs and intestines removed) of flies were dissected in PBS and fixed in 4% paraformaldehyde for 20 min, then washed twice with PBS, incubated for 20 min in fresh Oil Red O solution (6 mL of 0.1% Oil Red O in isopropanol and 4 mL distilled water, and passed through a 0.45 mm syringe), followed by rinsing with distilled water. Bright-field images were collected using a Leica M165 FluoCombi stereoscope system (utilizing a single focal plane) and processed using Leica software and Adobe Photoshop. Note: Contrast (red – neutral lipids vs. yellow/black – cuticle) was enhanced using Adobe Photoshop (equal for all images) in order to better visualize the red stain.

### Oral infection assays

*Pseudomonas entomophila* (*P.e.*) was used for natural (oral) infections. Briefly, for oral infection, flies (7-10 d of age) were placed in a fly vial with food/bacteria solution and maintained at 25 °C. The food solution was obtained by mixing a pellet of an overnight culture of bacteria (OD_600_=50) with a solution of 5% sucrose (50/50) and added to a filter disk that completely covered the surface of standard fly medium. Fat body of infected flies were dissected 20 h after oral contact with infected food.

### Cell culture, plasmids construction and transient transfection

Drosophila S2R+ cells (obtained from Drosophila Genomics Resource Center, DGRC) were maintained in Schneider’s Drosophila media supplemented with 10% FBS, 50 U/mL penicillin, and 50 μg/mL streptomycin at 25 °C.

Relish coding sequence was amplified from pMT-Flag-RELISH (Addgene plasmid # 63750; http://n2t.net/addgene:63750; RRID: Addgene_63750), and then cloned in the pAC5.1 V5-His plasmid (pAC5.1-Relish-V5).

HDAC6 coding sequence was amplified from UGO01255 (DGRC Stock 1663924; https://dgrc.bio.indiana.edu//stock/1663924; RRID: DGRC_1663924), and then cloned in the pAC5.1 V5-His plasmid (pAC5.1-HDAC6-HA).

After sequencing, 1 μg of plasmids was transfected into S2R+ cells (2×10^6^) using Effecten transfection Kit (Qiagen).

### Lysis of cells and flies, and immunoprecipitations

Cells were rinsed with cold phosphate-buffered saline (PBS) and lysed in IP lysis buffer with 1x Protease Inhibitor cocktail (Thermo Scientific). Cell lysates were cleared by centrifugation in a microcentrifuge (15,000 x rpm for 10 min at 4 °C). Cell lysate samples were prepared by the addition of 4x Laemmli SDS sample buffer (Thermo Scientific), resolved by 8–12% SDS–polyacrylamide gel electrophoresis (PAGE), and analyzed by immunoblotting.

Whole flies were crushed physically using a bead beater in IP lysis buffer with 1x Protease Inhibitor cocktail (Thermo Scientific). The resulting lysates were cleared by centrifugation in a microcentrifuge (15,000 x rpm for 10 min at 4 °C) and analyzed as above.

For anti-V5 immunoprecipitations, 500 μL cleared lysates were incubated with 1 μL Mouse anti-V5 antibody (Sigma, V8012) overnight at 4 °C. Magnetic Protein A/G beads (Pierce) were washed three times with IP lysis buffer. A 30 μL volume of resuspended beads in lysis buffer was added to cleared lysates and incubated overnight at 4 °C. Immunoprecipitates were washed five times; three times with IP lysis buffer and twice with IP lysis buffer with 500 mM NaCl. Immunoprecipitated proteins were denatured by addition of 50 μL of SDS sample buffer and heated at 100 °C for 5 min. Denatured samples were resolved by 8–12% SDS–PAGE, and analyzed by immunoblotting.

### Immunoblotting analysis

Fat body/carcasses from 10 flies (per sample) were dissected in PBS and then homogenized in a protein sample buffer. Experiments with the separation of nuclear and cytoplasmic proteins were performed using a Nuclear and Cytoplasmic Protein Extraction Kit (Invent Biotechnologies, NT-032) according to the manufacturer’s instructions. Proteins were separated by SDS-PAGE and transferred to PVDF membrane using standard procedures. The following antibodies were used: Rabbit anti-HA (Cell signaling technology, 3724S, 1:1,000), Rabbit anti-beta-actin (Cell signaling technology, 4967S, 1:1,000), Rabbit anti-Relish (Raybiotech, RB-14-0004, 1:5,000), Rabbit anti-V5 (Sigma, V8137, 1:5,000), Rabbit anti-Histone H3 (ABclonal, A2348, 1:5,000), Mouse anti-GAPDH (ABclonal, AC002, 1:2,000), and Mouse anti-Flag (Sigma, F1804, 1:5,000) were incubated with the membrane overnight at 4 °C. Secondary anti-mouse (BIO-RAD, 1:10,000) or anti-rabbit (BIO-RAD, 1:10,000) was incubated for 1 hr at room temperature. Signal was detected using HRP-conjugated anti-rabbit and ECL Western Blotting Substrate (Pierce), according to manufacturer instructions.

### Quantification and statistical analysis

Samples sizes were predetermined by the requirement for statistical analysis, and at least 3 biological replicates were utilized for all experiments. For all quantifications, n represents the number of biological replicates, and error bar represents SE or SEM. Statistical significance was determined using either the unpaired *Student’s t*-test in GraphPad Prism Software, and expressed as p values. (*) denotes values whose difference was significant, and (n.s.) denotes values whose difference was not significant.

## Data availability

Data and materials that support the findings are available from the corresponding author on request. All relevant data supporting the findings of this study are available within the article and its Supplementary Information files.

## Acknowledgments

This work was supported by the National Institute of Diabetes and Digestive and Kidney Diseases (grant R01 DK133294 and grant R56 DK108930 to J.K.).

## Author contributions

X.K. designed and performed experiments, as well as wrote the manuscript. C.L. performed experiments, J.K. designed experiments and wrote the manuscript.

