## Supplementary Information for "NF-κB restrains nutrient-dependent transcription programs through chromatin modulation in *Drosophila*"

**Table S1: qRT-PCR Primers**

| Primer | Sequence (5’-3’) |
|---|---|
| actin5C-F | CTCGCCACTTGCGTTTACAGT |
| actin5C-R | TCCATATCGTCCCAGTTGGTC |
| Lamp1-F | ACCTATGAGGCTAGGGAGGG |
| Lamp1-R | GGAAAGCACAGTGGTGTCAT |
| Atg1-F | CACGGCACACATTCTGCTTC |
| Atg1-R | CAACGGCATCGCGATATTTGT |
| Rab40-F | GCGAGTCCTTTTGCGAACTG |
| Rab40-R | CCTGAAGCGACAGCACCTTA |
| Bmm-F | CAATAAGGGTCTGGCCAACTGGAT |
| Bmm-R | TAAGTCCTCCACCATTACTCTGGC |
| CaMKII-F | TTCACCATCGTTTTCCTGCT |
| CaMKII -R | GGCAGCTTGAGTCAGGTGAT |
| mdy-F | CGGAACTCCCTGATTCCGTT |
| mdy-R | AGATGATTGGGTAGCGCAAGT |
| ImpL2-F | AACCGCGAATCATCTACACC |
| ImpL2-R | GGACGATCTCCTTGTTCTCG |
| wdb-F | GACGTGTTCCCAGAAGGAGG |
| wdb-R | GCTCGCTCAGCAACCTGAAA |
| trbl-F | CGAGGGCCATATTCCACCAG |
| trbl-R | TACTGCAGTTTCGTTCTGGC |
| Itpr-F | AGCCAAGGAAGTAGCACCAC |
| Itpr-R | ACAAACAAGCCCTCCTTGCT |
| yellow-h-F | ATCCGGATCCCTTTGCTTCG |
| yellow-h-R | AGGCACCATGAACTCCTTGAA |
| Hgo-F | TTACCGTGCTCACTTGTCCC |
| Hgo-R | GCTCATGCAGTTCCTGTGGT |
| trh-F | CACTGCGAGCCAAGGGTATC |
| trh-R | CGATCAGGTCTGAGTGGCTC |
| Hpd-F | ATTCGTGAGATTCGCTACCATC |

| | |
|---|---|
| Hpd-R | GGCAATCCGTTTAGAAGGACATC |
| Tat-F | TGATTGATGTGCTCCACTCGAA |
| Tat-R | TGGACAAAGTGGGTGTCGTC |
| Faa-F | TTCCCTATCTGCGCCAGAAC |
| Faa-R | GTTTTGATCGGCGGGTTTCAA |
| Hydr2-F | TCAAGGAGTATGCCACGACG |
| Hydr2-R | GCATCATGACCAAGGACCCA |
| Oapt74D-F | GCTTGCATTCCAGATGCTCC |
| Oapt74D-R | GCCGACTGTATCACACCCAT |
| Pepck1-F | ACAAAGCAAAATTATCTCCGGCA |
| Pepck1-R | AGATGGGAATGCCACGGATG |
| Pepck2-F | TGTCATTCCTTTTAGCATGGGTC |
| Pepck2-R | GACGACGTATGGTGAGTCTGT |
| Sik3-F | CGCGATCAAATGATCCCCCT |
| Sik3-R | GTAGGGAACTGGCTGAGTGG |
| Glys-F | CGCACCCTGGGCATTTACTA |
| Glys-R | GAGTCCTCGTCATCCGAACC |
| Glyp-F | CCACTACTACTTGCTGGCCG |
| Glyp-R | CCACTTGGCTTGGTTCTGGT |
| Dg-F | GCAAAATGGAGCTAGGCGAC |
| Dg-R | TTGTAGGATGGTGGCAGCAG |
| ActinBeta-F | TGTGGTTGCCCTTAGCCTTG |
| ActinBeta-R | ATTGGCGCTCAGTTAGGACA |
| Hsp83-F | CCCTTACTGACCCCAGCAAG |
| Hsp83-R | GATGGTCAGAGTACCAGCCG |
| Usp47-F | CATCGCAGCGGAGTTCATTG |
| Usp47-R | CGGACACCACCGTCTTACAA |
| Cdc37-F | TTCGACAGCGAGATCGAAGG |
| Cdc37-R | AGGCCTTTAGTTCATCGGGC |
| step-F | GGATAAGGAACCACGCGGAA |
| step-R | TTGCCCTCTACCACTTTGCC |

| | |
|---|---|
| melt-F | TACTCAAGTGCGAGAACCGC |
| melt-R | GGATCTCGCGGTCCTTAACC |
| CycG-F | TCGTTCCACTTGGCCATCAA |
| CycG-R | AGCGGTACAACCACACTGAG |
| Lin28-F | CCGCACAAGAATGTGACCCA |
| Lin28-R | TCTTCTCTTGGGCTGCACTG |
| Gbs-76A-F | ATGTCCTCATGTCGCGGAAG |
| Gbs-76A-R | CCTCCTCGGTTGTCTGTGAT |
| Hr96-F | TTGGATCGCGAGCTAAACGA |
| Hr96-R | CGAGTGTCGTCGGGCTTAAT |
| Gbs-70E-F | ATCCTCCGGATCAGTGGACA |
| Gbs-70E-R | AGTCGCGTTCTATCTTTTGTCTAA |
| Pask-F | TGCGTTAATGAAGCCAAGCAC |
| Pask-R | GACCCTCAGCTCGTGGTTAC |

**Table S2: ChIP-qPCR Primers**

| Primer | Sequence (5'-3') |
|---|---|
| ChIP-Atg1-F | GAAAGTGGTATTTTTCGCGC |
| ChIP-Atg1-R | ACGATCAGCGTAATTTCCTT |
| ChIP-BmmR1-F | GCTTGTTTGCGTTTGTAGGTC |
| ChIP-BmmR1-R | TTCGAAATTGGACAAACACG |
| ChIP-BmmR2-F | TGTCGCTGACAATCAAAAGC |
| ChIP-BmmR2-R | TTCTGGGTGGAGTTTGGAAC |
| ChIP-Faa-F | AAGTTCTGTGGTGGTTTCCG |
| ChIP-Faa-R | TATGGGTTCACCCAGCTGAT |
| ChIP-Tat-F | TAAAGGGCTCCAAGCTATCC |
| ChIP-Tat-R | TTTATGGTGCTCACCAATGG |
| ChIP-Hpd-F | GGAGATTTCCTGCCTGGATT |
| ChIP-Hpd-R | GCGGAGTATTCCGTGTGAAT |

### Supplementary Figures

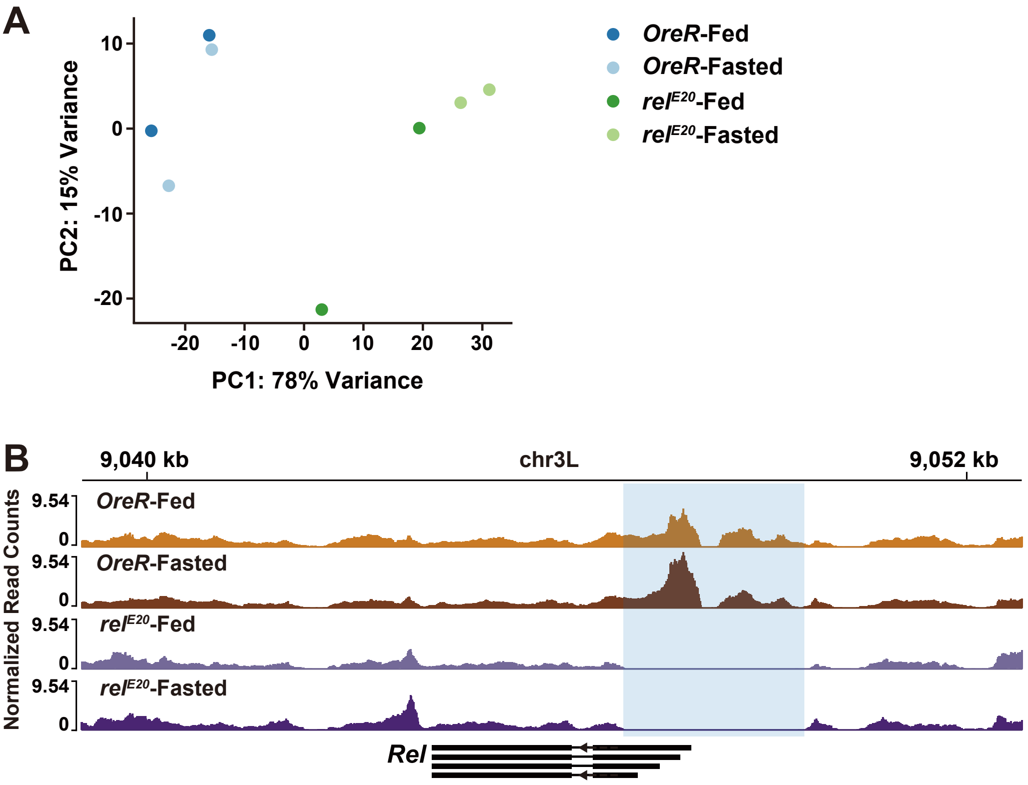


**Figure S1. Landscape of DNA accessibility in *OreR* control and Relish mutant flies. Related to Figure 1.**

(A) PCA of peak accessibility in all samples. Each symbol represents an independent biological replicate. PC1 (78%) and PC2 (15%). (B) Normalized chromatin accessibility profiles at *Rel* gene locus. Shaded region indicates the genomic region deleted in Relish mutant flies.

Fasted represents 20h of nutrient deprivation. All flies were 7 days old post-eclosion.

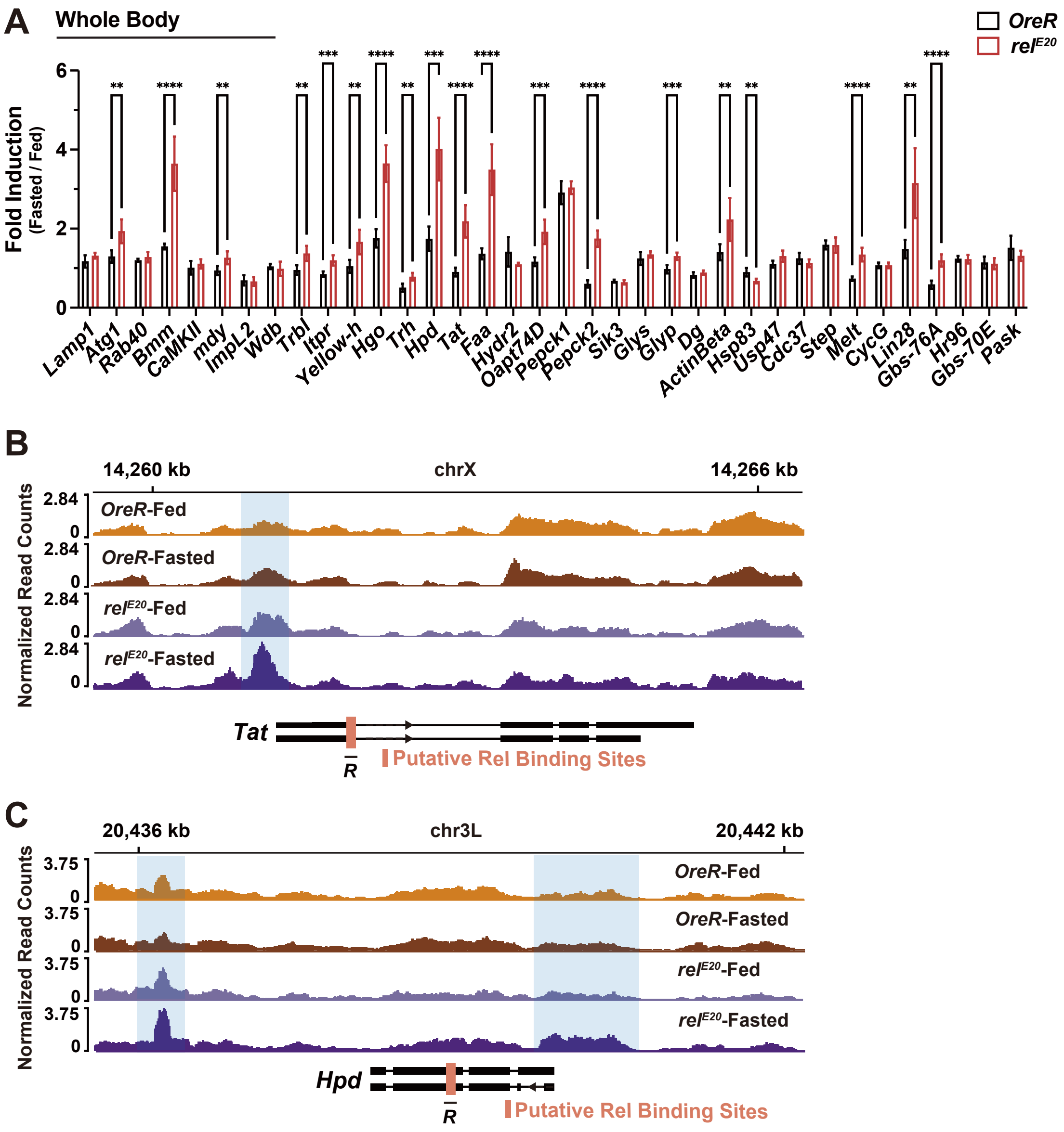


**Figure S2. Relish-dependent changes in metabolic gene expression and chromatin accessibility. Related to Figure 2.**

(A) Transcriptional changes (measured by qRT-PCR in the whole body of *OreR* control and Relish mutant flies, plotted as fold induction [ratio of 20 h fasted to fed] of relative expression) of distinct upregulated metabolic genes identified in Figure 1F. n = 5 replicates. (B and C) Normalized chromatin accessibility profiles at individual targeted gene loci. Shaded regions indicate chromatin peaks exhibiting uniquely increased accessibility in fasted Relish mutant flies. Red bars represent Relish binding sites predicted by JASPAR (relative profile score threshold > 75%). $\bar{R}$ represent regional target sites (and corresponding primer sets) tested in ChIP-qPCR analysis.

Bars in (A) represent mean ± SE. **p value < 0.01; ***p value < 0.001; ****p value < 0.0001. All flies were 7 days old post-eclosion.

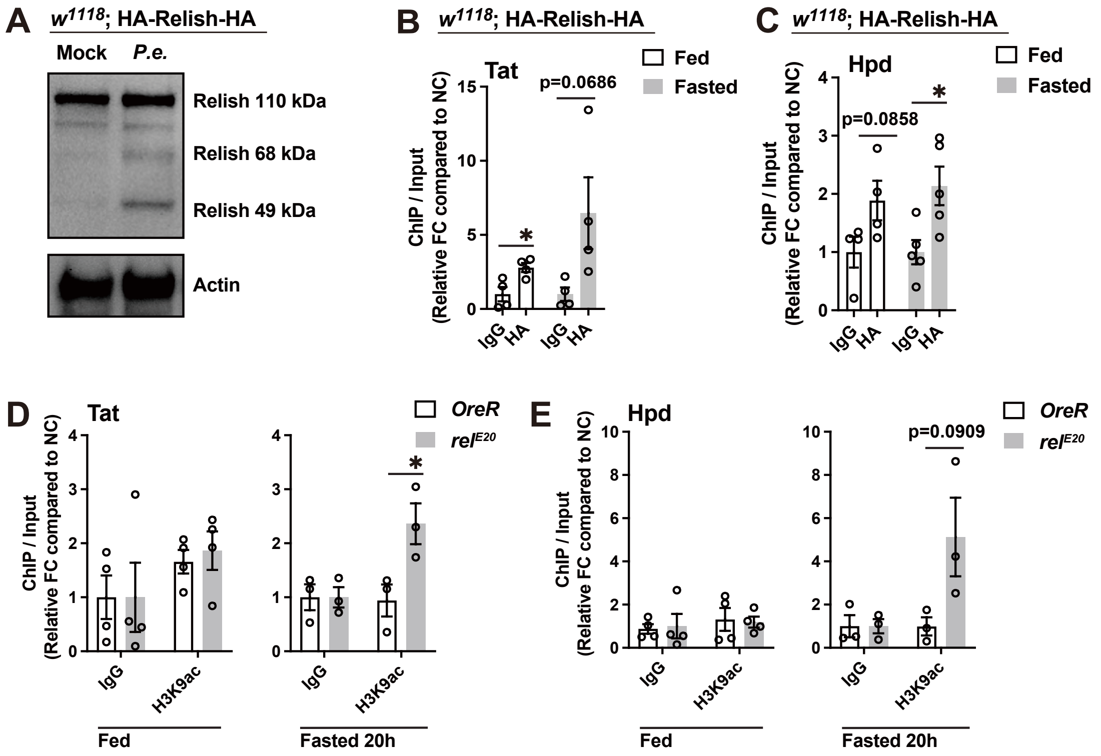


**Figure S3. Relish-dependent changes in chromatin modifications. Related to Figure 2.**

(A) The Relish protein and its cleaved forms upon bacterial infection. Genotype: *w1118*; HA-Relish-HA. (B and C) ChIP-qPCR analysis of Relish binding to the predicted sites in fed and fasted (20h) conditions (genotype: *w1118*; HA-Relish-HA). ChIP-qPCR analysis with normal IgG is included as a control. Plotted as fold change (FC) of indicated PCR primer sets compared to a negative control (NC) primer set. n= 4-5 biological replicates. (D and E) ChIP-PCR analysis of H3K9ac enrichment in Relish-binding regions in *OreR* control and Relish mutant genotypes before and after fasting (20 h). n = 3-4 biological replicates.

Bars in (B-E) represent mean ± SEM. *p value < 0.05. All flies were 7 days old post-eclosion.

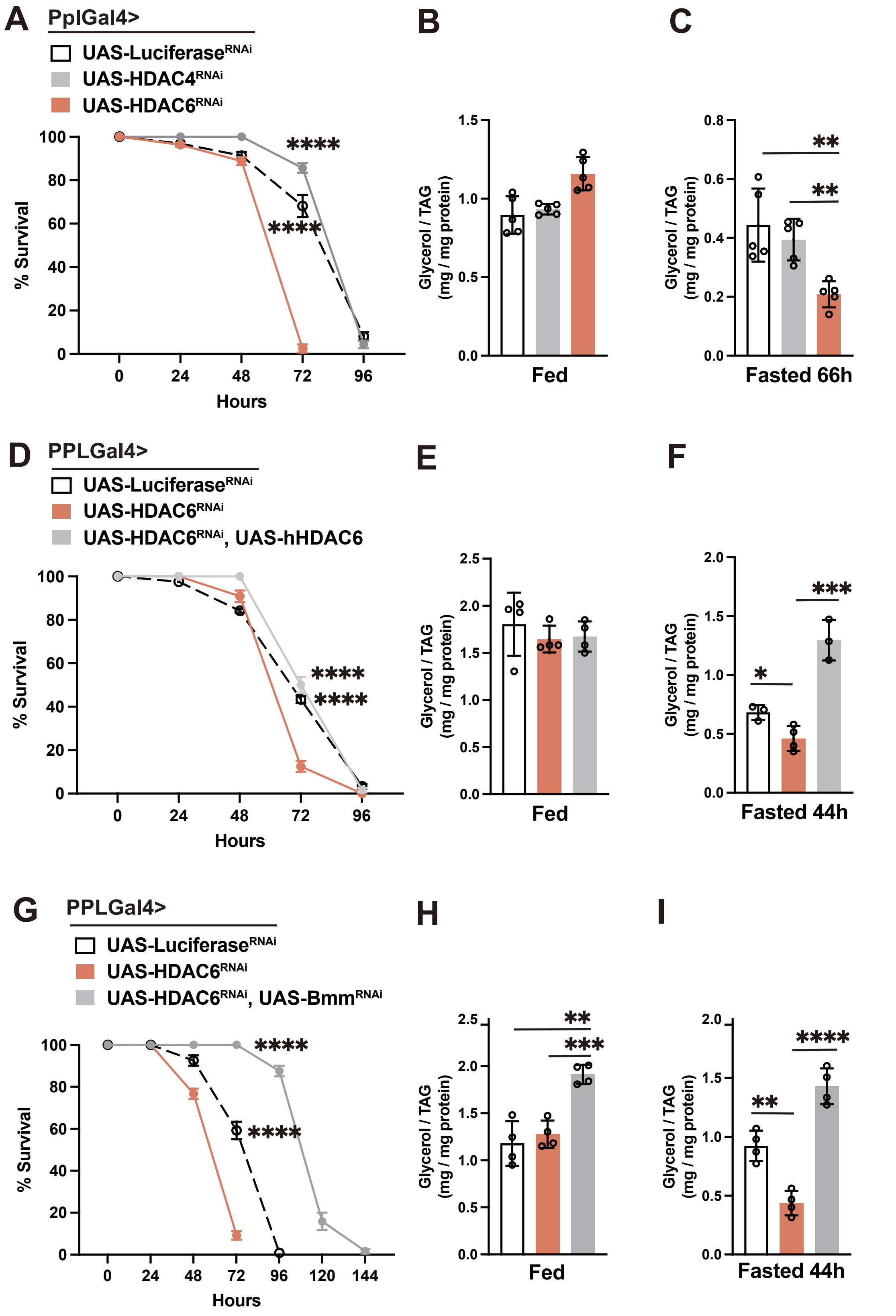


**Figure S4. HDAC6-dependent changes in lipid metabolism in response to metabolic adaptation. Related to Figure 4 and 5.**

(A) Starvation resistance of female flies (PplGal4/+; UAS-Luciferase$^{RNAi}$,

PplGal4/+; UAS-HDAC4$^{RNAi}$ or PplGal4/+; UAS-HDAC6$^{RNAi}$). n=8 cohorts (total 160 flies). (B and C) Total TAG levels of whole bodies during feeding and fasting conditions (n = 5 samples). (D-F) Re-expressing hHDAC6 in the fat body of HDAC6-deficient flies restores metabolic adaptation responses. (D) Starvation resistance of female flies (PplGal4/+; UAS-Luciferase$^{RNAi}$, PplGal4/+; UAS-HDAC6$^{RNAi}$ or PplGal4/UAS-hHDAC6; UAS-HDAC6$^{RNAi}$/+). n=6 cohorts (total 120 flies). (E and F) Total TAG levels of whole bodies during feeding and fasting conditions (n = 3-4 samples). (G-I) Attenuating Bmm in the fat body of HDAC6-deficient flies restores metabolic adaptation responses. (G) Starvation resistance of female flies (PplGal4/+; UAS-Luciferase$^{RNAi}$, PplGal4/+; UAS-HDAC6$^{RNAi}$ or PplGal4/+; UAS-HDAC6$^{RNAi}$/UAS-Bmm$^{RNAi}$). n=6 cohorts (total 120 flies). (H and I) Total TAG levels of whole bodies during feeding and fasting conditions (n = 3-4 samples).

Bars in (A, D and G) represent mean ± SEM, bars in (B, C, E, F, H and I) represent mean ± SE. *p value < 0.05; **p value < 0.01; ***p value < 0.001; ****p value < 0.0001. All flies were 7 days old post-eclosion.

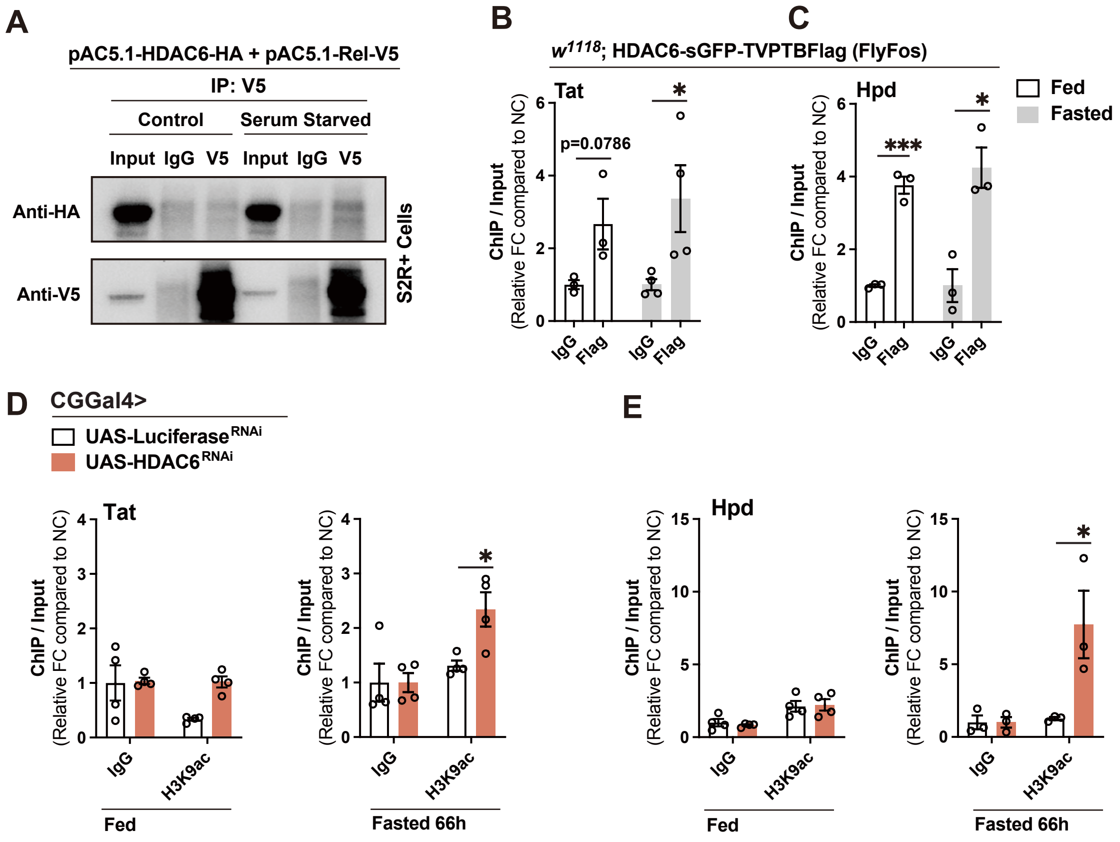


**Figure S5. HDAC6-dependent changes in chromatin organization in response to metabolic adaptation. Related to Figure 6.**

(A) *In vitro* interaction of Relish with HDAC6. Immunoprecipitates were prepared from lysates of S2R+ cells transfected with plasmids pAC5.1-HDAC6-HA and pAC5.1-Rel-V5. (B and C) ChIP-qPCR analysis of HDAC6 binding to the Relish-binding sites in fed and fasted (20h) conditions (genotype: *w1118*; HDAC6-sGFP-TVPTBFlag). n= 3-4 biological replicates. (D and E) ChIP-qPCR analysis of H3K9ac enrichment in Relish-binding regions in CGGal4/+; UAS-Luciferase$^{RNAi}$ and CGGal4/+; UAS-HDAC6$^{RNAi}$ genotypes before and after fasting (66 h). n = 3-5 biological replicates.

Bars in (B-E) represent mean ± SEM. *p value < 0.05; ***p value < 0.001. All flies were 7 days old post-eclosion.
